# Tuning siRNA packing order in lipid nanoparticles modulates oligonucleotide functional delivery

**DOI:** 10.64898/2026.02.06.704289

**Authors:** Artu Breuer, Georgios Kyriakakis, Marcus W. Dreisler, Frank H. Schulz, Georgios Bolis, Stavroula Margaritaki, Vasilis Papageorgiou, Ntaniela Spacho, Nikos S Hatzakis

## Abstract

Efficient siRNA delivery by lipid nanoparticles (LNPs) is widely attributed to carrier composition, yet how intraparticle packing governs function remains unclear. Here, we developed a single-particle fluorescence microscopy assay that simultaneously quantifies size and siRNA loading of individual, chromophore-labeled LNPs. Imaging ~0.5M particles per condition per hour uncovered two major packing modes: a high and a low order corroborated by cryo-EM. Quantitative live cell imaging combined with systematic variation of LNPs lipid composition and N/P ratio allowed deconvolution of siRNA packing, internalization, and silencing and its dependance on lipid composition and electrostatics. Surprisingly, low-order particles while encapsulating modest RNA mediate more efficient knockdown of a fluorescent reporter than their high-order counterparts. Guided by these findings we predicted and experimentally validated that tuning composition and N/P ratio to favor less compact siRNA packing enhances silencing potency. This framework offers actionable guidance for the rational optimization of LNP formulations for RNA therapeutics.

## Main

Lpid nanoparticles (LNPs) have revolutionized nucleic-acid therapeutics, enabling systemic delivery of mRNA and siRNA, leading to the first approved mRNA vaccines^1–3^. Their clinical success arises from a modular composition in which ionizable lipids govern cargo encapsulation and pH-triggered membrane destabilization, phospholipids stabilize the particle scaffold, sterols tune membrane packing and fluidity, and PEG-lipids suppress aggregation while extending circulation time ^2^. Additional functionalization strategies, including antibody conjugation and SORT-like approaches, enable cell-specific^3^, and extrahepatic delivery^4^. Yet, the same compositional diversity that supports adaptability across RNA modalities also complicates mechanistic interpretation. Despite this progress, the physicochemical determinants of efficient cytosolic delivery remain elusive.

Intracellular RNA delivery is restricted by multiple barriers. Only ~1-5% of siRNA is calculated to escape into the cytosol, with release confined to a narrow endosomal window, showing escape as the principal bottleneck^5–9^. Time-resolved single-particle assays at endosomal membrane mimics reveal rapid acidification-driven disintegration events with heterogeneous, often partial cargo release, pointing to kinetic barriers beyond fusion^10^. Roughly 70% of internalized cargo is exocytosed, decoupling uptake from functional delivery^11^, and single-cell multiomic profiling reveals that cell-to-cell heterogeneity further modulates LNP uptake and translation, compounding the difficulty of predicting delivery outcomes from ensemble measurements^12^. Membrane-damage markers such as galectin recruitment indicate endosomal rupture, but ESCRT-mediated repair rapidly reseals lesions^13^, so rupture alone is not a reliable proxy for productive release. Thus, while endosomal rupture is necessary, it may not be sufficient for cytosolic nucleic-acid delivery, indicating that particle-intrinsic structural features, remain poorly understood, could determine the functional outcome.

The internal mesoscopic architecture of LNPs, defined by lipid composition, cargo identity, and N/P ratio, is intrinsically heterogeneous. Cryo-EM and SAXS reveal composition-dependent mesophase behavior^10,14–18^, often modulated by N/P ratio: mRNA LNPs adopt pH-triggered inverse and non-lamellar phases^15^, whereas siRNA LNPs remain largely lamellar, with RNA localized between apposed bilayers surrounding an amorphous lipid core^16,19,20^. The identity and molar fraction of the ionizable lipid, together with the N/P ratio, govern this internal order and balance encapsulation efficiency, membrane integrity, and potency^14,17^. Correlations between multilamellar-to-amorphous transitions, uptake and silencing have been reported, yet these primarily relied on photoactivation to induce endosomal release^21^ and provided clues on how lipid-siRNA organization regulates LNP responsiveness.

Phase organization is reported to influence delivery efficiency^22–28^. Conventional techniques such as dynamic light scattering and bulk fluorescence assays while convenient, provide only population-averaged readouts of LNP size and loading. These mask due to ensemble averaging distinct subpopulations that differ in morphology and composition and thereby obscuring the nanoscale heterogeneity underlying functional diversity^23–26,29^. Single particle studies offer a route to address this intrinsic heterogeneity. Li et al.^27^ quantified size-payload variability by single-particle spectroscopic chromatography, revealing pronounced compositional dispersion yet without linking it to delivery efficacy. Kamanzi et al. combined multiparametric CLiC microscopy with alternating laser excitation to simultaneously measure size, FRET-based structure, and mRNA payload of individual suspended LNPs, further highlighting formulation-dependent heterogeneity but without connecting these properties to functional delivery^30^. Cryo-EM studies showcased that inverse siRNA hexagonal phase enhances the intracellular silencing efficiency as compared to multilamellar phases^22^, and Sjöberg et al.^28^, combining scattering and fluorescence microscopy on supported lipid bilayers, linked LNPs structural heterogeneity to loading and fusion. While these studies highlight the potential of single-particle methods to disentangle heterogeneity in morphology, loading, and dimensions, they do not provide direct links of single-particle heterogeneity to functional delivery.

Here, we developed a high-throughput single-particle fluorescence assay that simultaneously quantifies diameter and siRNA copy number for chromophore-labeled lipid nanoparticles (>0.5×10^6^ particles per condition in less than 2 hours). This multimodal framework reveals two general packing orders, low-order and multilamellar high-order, validated structurally by cryo-EM. Quantitative live-cell imaging of nanoparticle uptake and gene silencing quantified how packing mode modulates siRNA functional delivery. Systematic tuning of lipid composition and N/P ratio shifts the balance between the two packing states. Counterintuitively, low-order particles encapsulate modest amounts of siRNA yet mediate efficient cytosolic release and target knockdown, whereas high-order, cargo-rich particles remain inactive. By linking nanoscale organization to functional delivery at the single-particle level, this framework transforms LNP packing order into an actionable design parameter for RNA therapeutics.

### Single-particle assay for size and cargo quantification

We established a generalizable single-particle fluorescence assay utilising total internal reflection fluorescence (TIRF) microscopy to synchronously extract both the nanoscale dimension of each LNPs and the copy numbers of its nucleic acid cargo. The methodology, adapted from our earlier work on liposomes^31–34^, integrates Gaussian-based particle analysis and calibration against DLS, and quantifies the dimensions of each diffraction-limited LNP without the need of super-resolution microcopy (Fig. 1a). Besides circumventing ensemble averaging^4,5^ and operating without the need of specialized instrumentation^6–8^ it enables analysis of >10^5^ particles per condition (~10^6^ across replicates Supplementary Fig. S1). ^6–8^

**Fig. 1:**
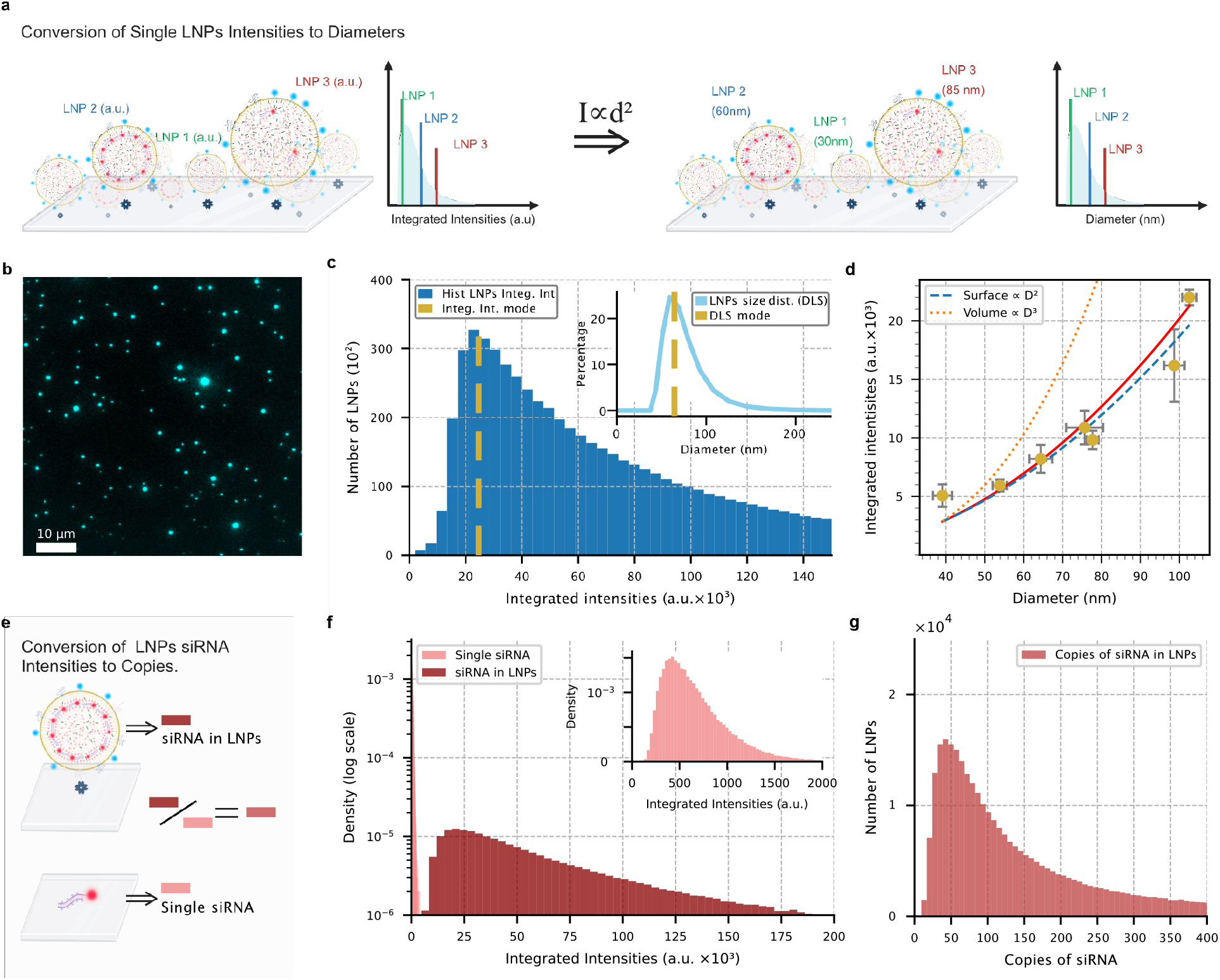
Conversion of fluorescence signals into single-LNP size and cargo content. **a**, Schematic illustrating the conversion of fluorescence signal from fluorescently labeled lipids into LNP diameter, based on the scaling of fluorescence signal with the square of the particle diameter (d^2^). **b**, Fluorescence micrograph showing individual LNPs tethered on passivated surfaces using streptavidin biotin method. LNPs are below the diffraction limit and appear as point spread functions (PSFs). Notice the large heterogeneity in signal. **c**, Histogram of background corrected integrated fluorescence intensities from ~10^6^ individual LNPs fluorescently labeled with DPPE-ATTO488 (dark blue), with mode intensity indicated (gold). Inset, corresponding size distribution by number from DLS (light blue) with mode size (gold). **d**, Scaling of the mode of integrated lipid fluorescence intensity for seven LNP populations with identical composition and varying diameters, plotted as a function of the mode diameter measured by DLS (error bars St.dev from 3-4 technical replicates). Blue and yellow dotted line correspond to modelled integrated intensity scaling with a cubic or quadratic relationship respectively **e**, Schematic illustrating the conversion of background corrected fluorescence of labeled siRNA to copy number of siRNA encapsulated in each individual LNPs into siRNA. **f**, Histogram of integrated fluorescence intensities from individual LNPs in the siRNA channel (dark red) and of single labeled siRNA molecules (light red). Inset, zoom-in of the single-siRNA intensity histogram. **g**, Histogram of measured siRNA copy numbers per LNP, derived from panel f.

LNPs containing DPPE-Atto488 as a size-reporting lipid, and siRNA-AlexaFluor647 were immobilized on functionalized coverslips^35,36^ and imaged by TIRF (Fig 1b). While each LNP is a diffraction limited spot, its integrated intensity depends on particle dimensions. From these measurements, we extracted the integrated intensity of >10^5^ LNPs per condition, yielding apparent size distributions in arbitrary units^31–34^ (Fig. 1c). Because DLS reports the most frequent physical diameter of the LNPs population, i.e. the mode, and TIRF reports the mode integrated intensity of the same population, the two can be directly paired, providing a reference point to link single-particle fluorescence to physical size (Fig. 1c). However, this anchor alone is insufficient; one must also establish how fluorescently labeled lipid scales with particle size.

To determine the scaling relationship, we generated LNP populations with tunable diameters by varying microfluidic formulation parameters while maintaining constant composition (Supplementary Table S1), producing mode diameters ranging from ~40 to >100 nm, as measured by DLS (Supplementary Fig. S2). Plotting integrated DPPE-Atto488 fluorescence against DLS-derived diameters (Fig. 1c-d) revealed a quadratic relationship (I ~ D^2^), consistent with a surface distribution of the DPPE-Atto488 labels (Fig. 1d), as we have previously shown for liposomes^31–34^. This surface localization is consistent with the net negative charge of DPPE-Atto488 at both physiological and acidic pH, which prevents electrostatic attraction to the negatively charged RNA cargo during and after formulation. By contrast, positively charged lipid dyes can redistribute toward the RNA-rich core, decoupling fluorescence intensity from particle dimensions^30^. Together, these findings demonstrate that lipid-associated fluorescence provides a quantitative proxy for particle diameter with nanometer-level precision.

We next calibrated the cargo channel to quantify the number of siRNA molecules per LNPby comparing integrated siRNA-AF647 fluorescence with that of single surface-immobilized siRNA reference molecules (Fig. 1e). After correcting acquisition settings (Supplementary Fig. S3), the ratio provided a direct estimate of siRNA copy number per LNP, completing the framework for simultaneous, quantitative measurement of size and cargo at the single-particle level. The method provides absolute quantification of nanoparticle heterogeneity across >10^5^ particles per condition (Supplementary Fig. S1) and is broadly compatible with standard fluorescence microscopes.

### Heterogeneity in LNP size and RNA loading

Imaging LNPs in both lipid and siRNA channels revealed striking particle-to-particle heterogeneity in cargo content relative to size (Fig. 2a-c). A zoom-in of a representative region, rendered as a 3D surface plot, highlights this variability: particles 1 and 2 exhibit comparable lipid intensities, consistent with similar diameters, yet differ vastly in siRNA content. Whereas particle 3 is larger than particle 1 but carries fewer siRNA molecules. These examples (Fig. 2c), demonstrate that LNP size does not deterministically predict RNA cargo loading.

**Fig. 2:**
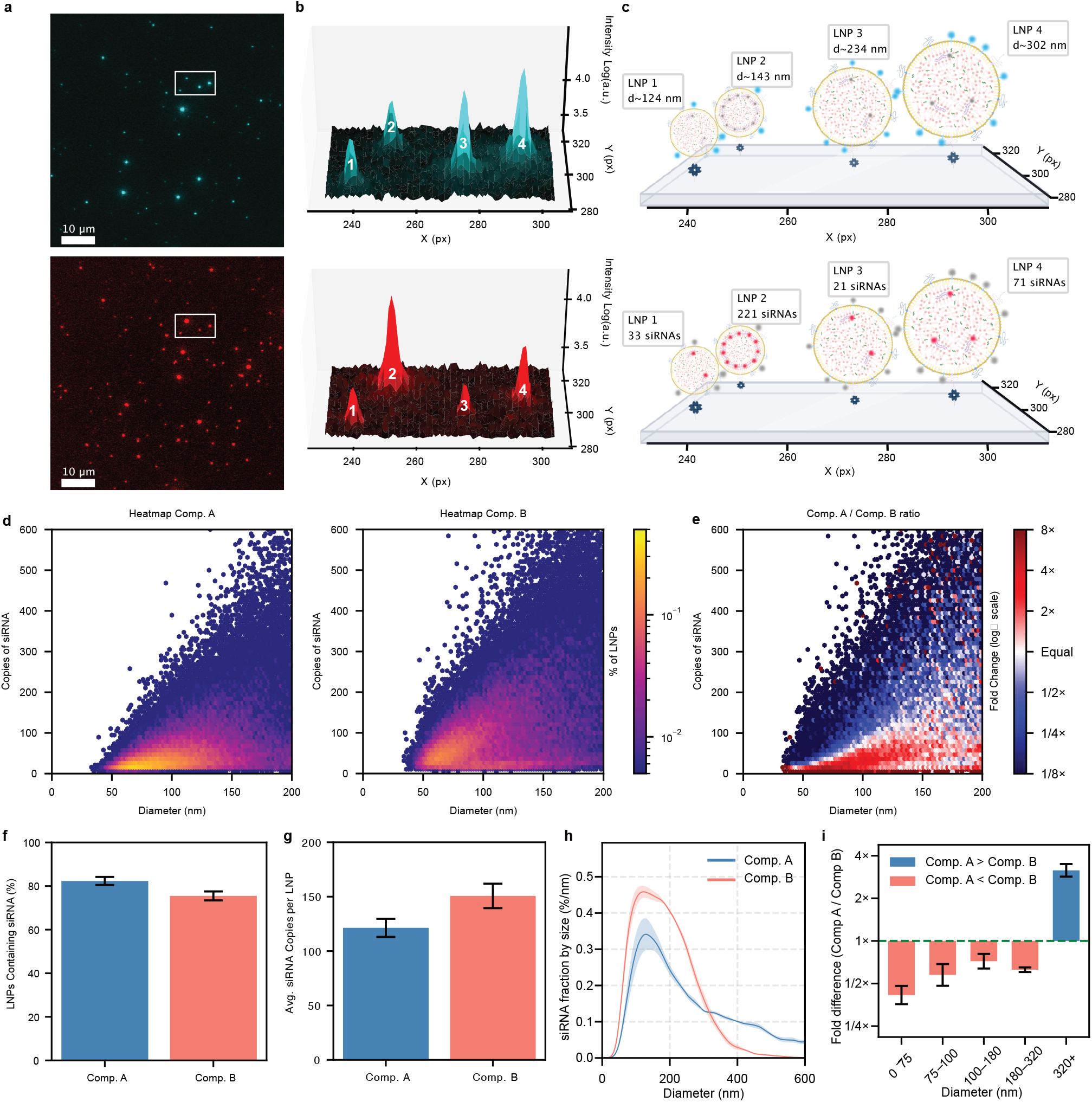
Quantification of single LNP heterogeneity in size and siRNA loading for two lipid formulations. **a**, Fluorescence TIRF micrographs of individual LNPs in the lipid (top) and siRNA (bottom) channels. **b**, 3D surface plots of fluorescence intensity within the white rectangle in a, showing LNPs in the lipid (top) and siRNA (bottom) channels. LNPs are heterogenous in dimensions and in siRNA loading. **c**, Schematic of the four LNPs shown in b, annotated with their estimated diameters (top) and siRNA copy numbers (bottom). Notice, the particle-to-particle heterogeneity in cargo content relative to LNP size. **d**, Heatmaps of siRNA copy number as a function of LNP diameter for lipid composition A (left) and composition B (right), based on >100,000 single LNPs per condition. Data granularity reveals that lipid composition affects loading **e**, Ratio metric heatmap showing the fold difference in LNP density between compositions A and B across the diameter-cargo space. **f**, Bar plot showing the percentage of LNPs containing siRNA for each composition, calculated over the entire population. **g**, Bar plot showing the average number of siRNA copies per LNP for each composition. **h**, Kernel-density-based distribution of total siRNA content as a function of LNP diameter for composition A and B, shown as mean ± St. dev. **i**, Fold difference in the percentage of siRNA copies across five defined size bins: 0-75 nm, 75-100 nm, 100-180 nm, 180-320nm and >300 nm. All data correspond to 3-4 technical replicates and error bars correspond to St.dev. of the 3-4 technical replicates.

We next scaled this analysis to two well-established formulations differing in ionizable lipid content and composition, Comp A and Comp B (see Methods). Heatmap scatter plots of siRNA copy number versus particle diameter revealed distinct loading landscapes (Fig. 2d). Although both formulations exhibited similar size distributions (Supplementary Table S2), their cargo organization differed markedly: Comp B showed broader dispersion of siRNA copy numbers across sizes, whereas Comp A was enriched in higher siRNA content across much of the diameter range. Ratiometric comparison (Fig. 2e) highlighted distinct regions of enrichment, with Comp B strongly biased toward higher cargo content across a broad size range, while Comp A contributed more to the low-loaded small-particle regime.

Despite these the single-particle differences, bulk summary statistics painted a deceptively similar picture: Both formulations contained a comparable percentage of RNA-loaded particles (Fig. 2f) and siRNA copy numbers (Fig. 2g). Because such ensemble averages obscure how cargo is distributed across the size spectrum, we quantified the total siRNA content as a continuous function of LNP diameter (Fig. 2h). Both LNP formulation concentrated siRNA within the intermediate diameter range, with broadly similar maxima. However, Comp A progressively shifted a substantial fraction of its total siRNA into the largest particles, creating a pronounced high-diameter bias absent in Comp B. Comp A consistently depleted siRNA from small and mid-sized particles relative to Comp B, with inversion only in the largest bin (Fig. 2i; Supplementary Note S1). Thus, despite similar mean loading, the formulations diverged sharply in cargo allocation. Single-particle analysis reveals distinct loading landscapes, reframing particle-to-particle heterogeneity from a confounding feature of ensemble assays into a measurable variable.

### Bimodal siRNA packing within single LNP formulations

To uncover the structural basis of these heterogeneous loading distributions, we analyzed the joint size-cargo kernel density estimate (KDE) contour plot at single-particle resolution (Fig 3a). Visual inspection revealed two preferential modes of siRNA loading, a low-density band and a higher-density branch diverging progressively with particle size, suggesting a binary-like organization of RNA packing (Fig. 3a). Control experiments with mixed labeled and unlabeled siRNA confirmed that the observed bimodality is not driven from fluorophore quenching (Supplementary Fig. S4).

**Fig. 3:**
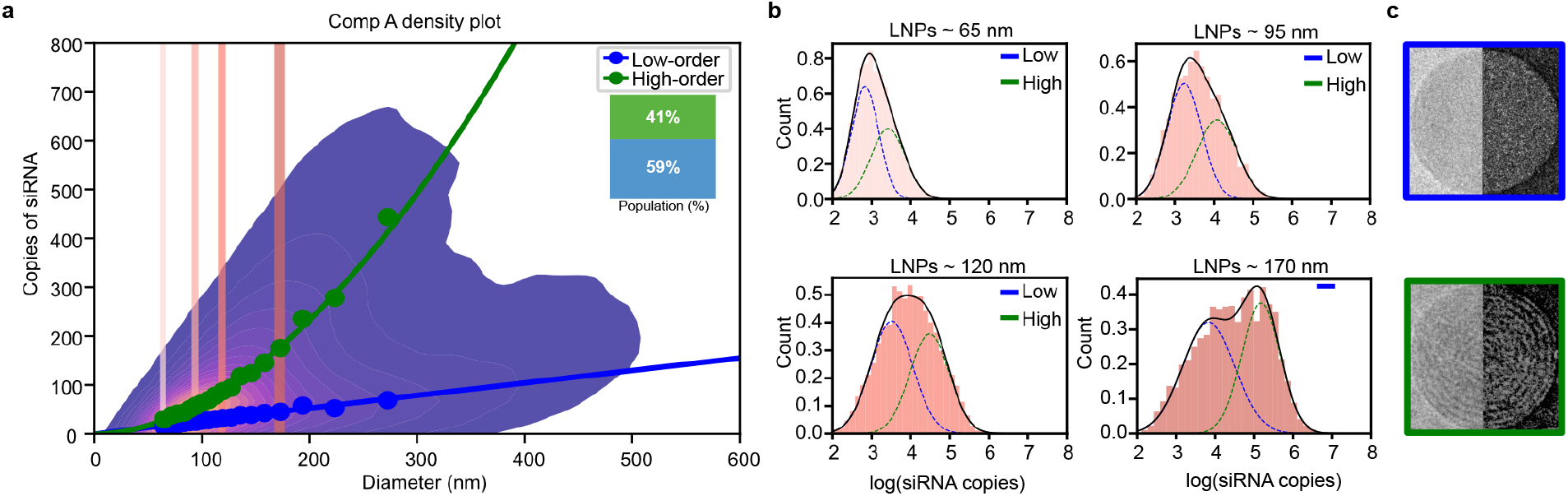
Heterogeneity in siRNA loading LNPs reveals distinct subpopulations with different scaling behaviors. Two-dimensional kernel density estimate (KDE) contour plot of siRNA copy number versus LNP diameter for formulation A reveals two distinct populations with different power-law scaling behaviors. Power-law fits are overlaid for the high-order (exponent: 1.92 ±0.08) and low-order (exponent: 1.05 ± 0.08) populations, averaged across four technical replicates. Inset indicates the relative abundance of each population (41±3% high-order and 59±3% low-order; n = 4 technical replicates). **b**, Log-transformed histograms of siRNA copy number for LNPs within four diameter percentile bins (55 nm, 85 nm, 115 nm, and 165 nm), corresponding to the 5th, 30th, 50th, and 75th percentiles, as indicated by color-matched tick marks in a. Two-component Gaussian mixture models reveal bimodal distributions corresponding to low- (blue) and high-order (green) loading states

To quantify this apparent bifurcation, we divided the size-cargo map into percentile-centered diameter bins and modeled the log-transformed siRNA distribution within each bin. The Bayesian Information Criterion (BIC) favored a two-component Gaussian model over one across most conditions (Supplementary Fig. S5), supporting low-load and high-load particle populations with diverse internal packing densities. Log-histograms from representative size bands confirmed the bimodal siRNA distributions with greater separation at larger diameters (Fig. 3b). Cryo-EM revealed two general morphologies: a low-density, low-order type corresponding to electron-lucent amorphous and bleb-like particles and a higher density, higher packing order type corresponding to multilamellar and locally inverse-hexagonal particles (Fig. 3c, Supplementary Fig. S6). This observation is consistent with the two populations identified by mixture modeling and earlier reports^16,19–21^, and verifies that the differential packing modes are not artifacts of chromophore quenching. Across the population, approximately 59±3% of particles belong to the low-order state and 41±3% to the high-order state (Fig. 3a), indicating that both architectures coexist in comparable proportions.

Plotting the component means in log-log space uncovered two separate scaling laws (Supplementary Fig. S7): slopes of 1.05 ± 0.08 for the low-order and 1.92 ± 0.08 for the high-order populations. In Fig. 3a, the corresponding power-law relationships are shown as blue and green lines overlaid on the linear-scale kernel density map, where their separation becomes visually evident. The weaker scaling of the low-order state is consistent with amorphous or disordered internal organization, whereas the near-quadratic dependence of the high-order state suggests multilamellar and locally inverse-hexagonal shell architectures, consistent with our and others Cryo-EM (Supplementary Fig. S7)^16,18,19,21,22,37^. Stochastic coagulation simulations reproduced comparable sub-volumetric exponents (Methods; Supplementary Fig. S8), supporting that the observed scaling arises naturally from probabilistic merger dynamics rather than surface-limited loading^38–40^. While the scaling exponents emerge from stochastic assembly, the coexistence of two distinct scaling regimes points to structural organization beyond stochastic variability: what appears as heterogeneity at the population level reflects two major self-assembled states with distinct physical scaling and functional potential.

These two packing states are consistent with earlier work on cryo-EM and small-angle X-ray scattering that correlated lipid composition and N/P ratio with lamellar, hexagonal, or inverted micellar arrangements^14,15,19,21^, and transitions from multilamellar to amorphous interiors with increasing N/P have been linked to LNP responsiveness to external stimuli and light-activated endosomal escape^21^. Notably, ensemble measurements of pH-triggered phase transitions capture population averages that necessarily obscure the coexistence of structurally distinct subpopulations^20,22^. The single-particle framework introduced here extends these insights in a critical respect: the two packing states are not points on a continuum but reproducible, discrete organizations whose relative abundance can be quantified across millions of particles and modulated by formulation conditions.

### Lipid composition tunes packing and functional delivery

To test whether lipid composition modulates RNA packing architecture, we compared Comp A and Comp B (See Methods, Supplementary Table S2). KDE plots for ~0.5 million particles per formulations revealed marked shifts in packing-state distribution (Fig. 4a,b), corroborated by cryo-EM micrographs (Supplementary Fig. S6). Comp A (N/P 3) contained 59 ± 3% low-order particles with a scaling exponent of 1.05 ± 0.08 (Fig. 4a), whereas Comp B (N/P 3) contained only 32 ± 1% low-order particles with a shallower exponent of 0.8 ± 0.2 (Fig. 4b). High-order scaled with exponents of 1.92 ± 0.07 (Comp A) and 1.32 ± 0.05 (Comp B), reproducible across 4 replicates (Supplementary Fig. S5, S9). Lipid composition thus reproducibly governs nanoscale packing order.

**Fig. 4:**
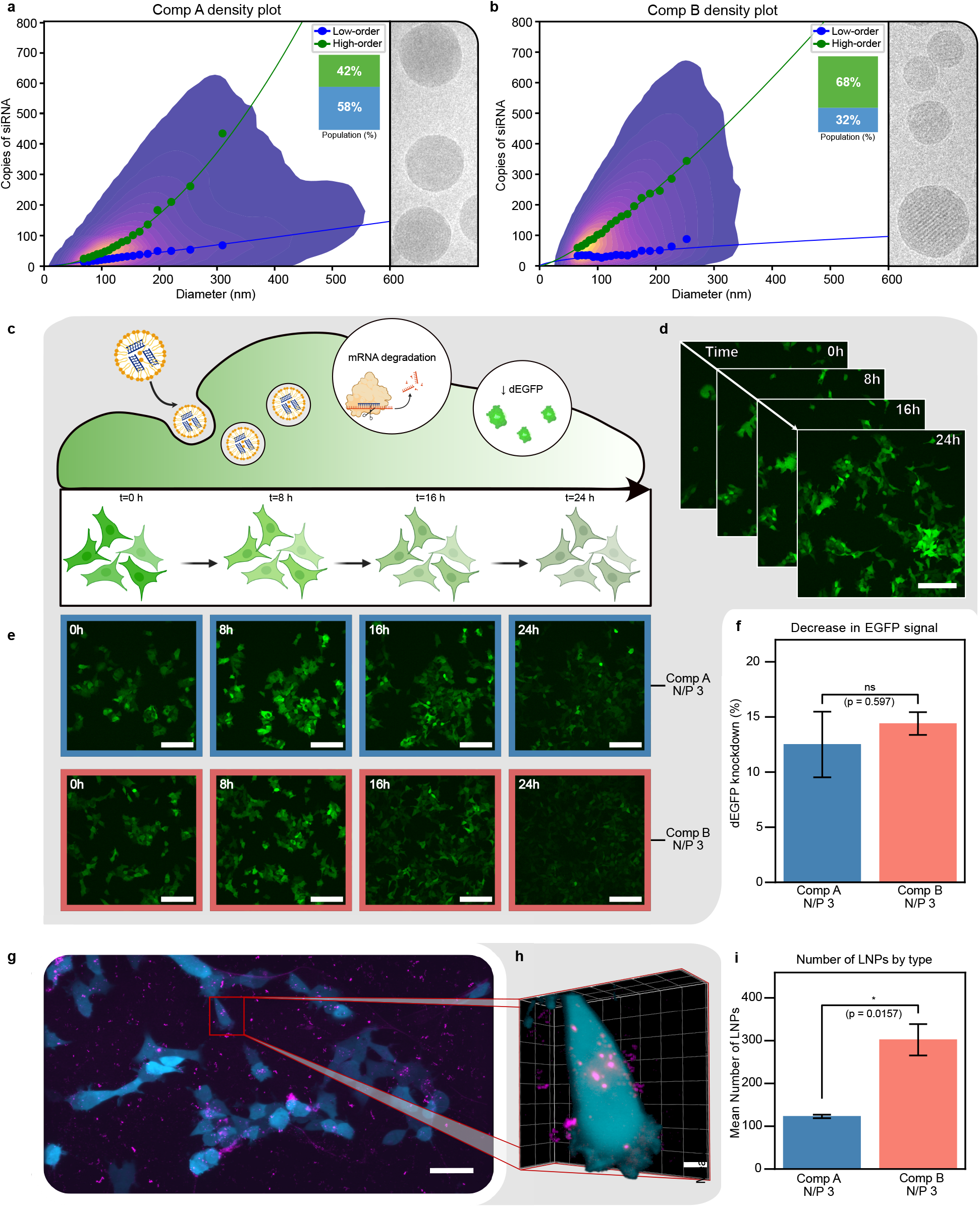
LNP composition modulates siRNA loading order influencing siRNA mediated target knockdown. **a, b**, Two-dimensional KDE contour plots of siRNA copy number versus LNP diameter for formulations A and B (left). KDE contour plots are representative of four independent technical replicates. Insets quantify the relative abundance of low- and high-order packing states, averaged across the four technical replicates. Representative cryo-electron microscopy images displaying mostly amorphous packing in A and mostly concentric architecture in B consistent with diverse packing of KDEs. **c**, Schematic illustration of experimental setup of cellular uptake of LNPs, siRNA release, and subsequent eGFP knockdown. **d**, Representative spinning-disk confocal time series of untreated HEK293T-d2eGFP cells (Scale bar, 50 μm). **e**, Representative spinning-disk confocal images of HEK293T cells expressing d2eGFP at 0 h, 8 h, 16 h, and 24 h following treatment with siRNA-loaded LNPs (300 nM). Images show the progressive decrease in eGFP fluorescence over the assay. For visualization, the z-plane with maximal eGFP fluorescence was selected, and display settings were kept constant across conditions (Scale bar, 50 μm). **f**, Quantification of d2eGFP knockdown after 24 h, showing reduced fluorescence intensity relative to untreated controls. Bars show mean ± S.E.M. from three independent technical replicates. Statistical significance was assessed using Welch’s t-test (P = 0.597; ns). **g**, Maximum-intensity projection of a lattice light-sheet microscopy (LLSM) dataset showing LNP internalization into d2eGFP-HEK293T cells (Scale bar, 50 μm). **h**, Representative 3D LLSM reconstruction of a single cell with internalized LNPs (Scale bar, 5 μm). **i**, Quantification of internalized LNPs per cell for formulations A and B. Bars show mean ± S.E.M. from three independent biological replicates. Statistical significance was assessed using Welch’s t-test (P = 0.0157; *).

We next examined whether packing order manifests in differences in uptake and functional delivery. Comp A and Comp B siRNA loaded LNPs (300 nM; Supplementary Table S3; Fig. 4c) were added to HEK293T cells stably expressing destabilized eGFP (d2eGFP). Time-lapse spinning-disk miscorscopy^41,42^ revealed progressive loss of fluorescence over 2h 4 (Fig. 4d,e; Supplementary Fig. S10). Quantification across three independent replicates confirmed comparable knockdown efficiencies of 12 ± 3 % for Comp A and 14 ± 1 % for B relative to untreated controls (Fig. 4f, Supplementary Fig. S11). However, lattice light sheet and quantitative image analysis revealed cells internalized ~2.5-fold more LNPs of Comp B (302 ± 37) than Comp A (123 ± 4) after 2.5 h of incubation (Fig. 4g-i; Supplementary Fig. S12), despite comparable hydrodynamic diameters (~80 nm; Supplementary Table S3 and Fig. S13), indicating that lipid chemistry rather than particle size alone, governs internalization efficiency^43^.

Interestingly if the amorphous fraction of each formulation is taken in consideration for internalization, <u>by multiplying cellular uptake by the amorphous fraction for each formulation, an</u> 1.3 ± 0.2-fold difference between formulation A and B emerges (73 ± 4 amorphous LNPs/cell for Comp A and 97 ± 12 for Comp B, respectively). This estimate was consistent with the measured fold difference in knockdown efficiency between formulations (Comp B/Comp A: 1.2 ± 0.3), suggesting that the effective dose of potentially release-competent particles may better explain functional delivery than total particle uptake alone. Mesoscale packing differences^44,45^, rather than the total amount of internalized RNA, thus appears to gate cytosolic release, with composition setting uptake and packing state determining the release-competent fraction. To assess whether packing order itself predicts release efficiency independent of lipid composition, we next modulated LNP self-assembly by varying the N/P ratio.

### Low-order packing drives productive siRNA delivery

To isolate the effect of packing order from lipid composition, we varied the N/P ratio while keeping formulation chemistry constant. KDE plots revealed a pronounced dependence of packing organization on N/P ratio (Fig. 5a-c). N/P 1 yielded exclusively high-order particles, whereas N/P 3 and 6 produced mixed populations low-order fractions of 59±3% and 61±2%, respectively, consistent with qualitative readout of Cryo-EM measurements (Supplementary Fig. S6). Condition-specific scaling exponents confirmed that N/P tuning alters both the relative abundance and the effective scaling of each packing mode: at N/P 3, low-order and high-order exponents were 1.05±0.08 and 1.92±0.08; at N/P 6, 0.85±0.10 and 1.43±0.17; N/P 1 displayed only the high-order branch (1.81 ± 0.07). Percentile-matched histograms confirmed unimodal intensity distribution at N/P 1 and bimodal mixtures at N/P 3 and 6 (Fig. 5d-f). These differences were reproducible across replicates (Supplementary Fig. S5, S14, S15), demonstrating that electrostatic stoichiometry directly governs nanoscale RNA packing.

**Fig. 5:**
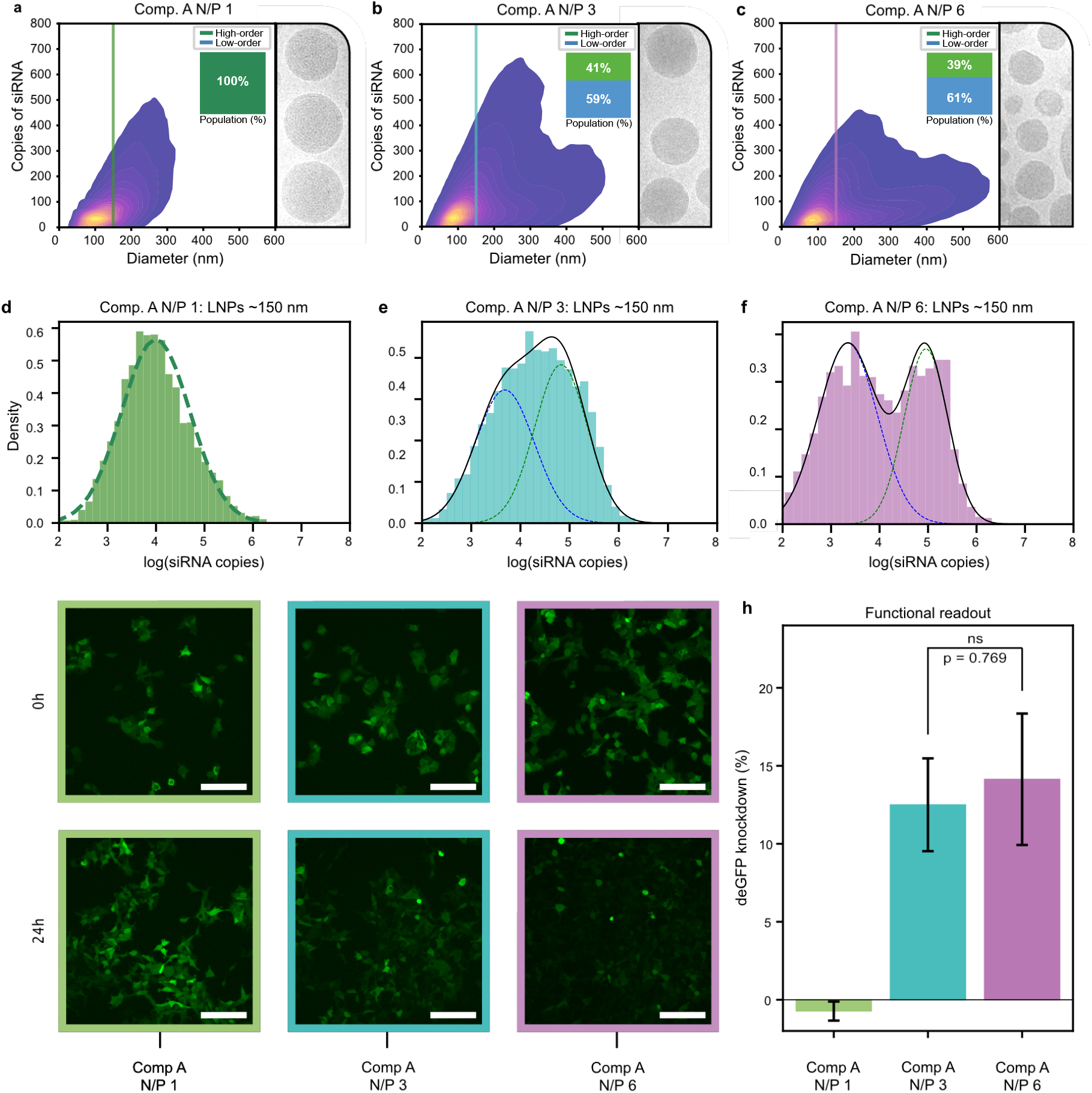
Low-order siRNA packing states correlate with enhanced siRNA-mediated knockdown activity. **a-c**, Effect N/P ratios to packing order. Two-dimensional KDE contour plots of siRNA copy number versus LNP diameter for formulation A at N/P ratios of 1, 3, and 6). KDE contour plots are representative of four independent technical replicates. Insets quantify the relative abundance of low- and high-order packing states, averaged across the four technical replicates. N/P 1 yields exclusively high-order particles, while N/P 3 and 6 show mixed populations. KDEs are consistent with representative cryo-electron microscopy images: **d-f**, Log-transformed histograms of siRNA copy number for LNPs within percentile bins centered around 150 nm, as indicated in a-c. Gaussian mixture modeling reveals unimodal distributions at N/P 1 (high-order only) and bimodal distributions at N/P 3 and 6, corresponding to low-order (blue) and high-order (green) subpopulations. **g**, Representative spinning-disk confocal images of HEK293-d2eGFP cells at the first and last time point of the 24 h knockdown assay (see Methods). Images illustrate the initial eGFP fluorescence level and its decrease following treatment. For visualization, the z-plane with maximal eGFP fluorescence was selected, and display settings were kept constant across all conditions (Scale bar, 50 μm). **h**, Quantification of d2eGFP knockdown in HEK293T cells after 24 h of treatment with siRNA-loaded LNPs (300 nM). Only formulations containing low-order subpopulations (N/P 3 and 6) induce knockdown. Purely high-order particles (N/P 1) display no measurable knockdown. Bars represent mean ± S.E.M. from three independent technical replicates. Statistical significance was assessed using Welch’s t-test; no significant difference was observed between N/P 3 and N/P 6 (P = 0.772; ns).

We next evaluated how these structural states influence functional delivery. Spinning-disk microscopy revealed progressive fluorescence decrease over 24 h following treatment with siRNA-loaded LNPs (300 nM; Supplementary Table S3; Fig. 5g; Supplementary Fig. S10). Quantification showed measurable knockdown for N/P 3 and 6 (12 ± 3% and 14 ± 4%, respectively, Fig. 5h), which were statistically indistinguishable (p = 0.772), consistent with their similar low-order fractions (59 ± 3% vs 61 ± 2%, p = 0.33) In contrast, N/P 1, lacking detectable low-order particles, produced no measurable knockdown, establishing that low-order, amorphous, architectures are essential for productive release, whereas purely high-order particles are functionally inert. Packing order, rather than lipid composition alone, thus governs siRNA delivery efficiency: electrostatic complexation not only dictates encapsulation but programs the internal order of the assembled nanoparticle, which in turn modulates functional outcome.

Mechanistically, these results suggest that low-order, amorphous particles represent deformable, metastable assemblies whose internal disorder facilitates cargo release. Amorphous organization may promote local membrane curvature fluctuations and expose ionizable lipid headgroups upon endosomal acidification, lowering the energetic barrier for fusion or transient pore formation^46^. Less densely packed siRNA within these particles may also be released more efficiently upon endosomal rupture than the tightly organized cargo in high-order counterparts. Higher N/P ratios increase the charge density at the point of complexation, promoting rapid charge neutralization and RNA condensation into compact, ordered assemblies. Conversely, near-stoichiometric conditions at lower N/P ratios yield more disordered assemblies, probably with greater water content and local structural defects, configurations that preserve encapsulation efficiency while remaining responsive to triggered disassembly. By contrast, stacked multilamellar bilayers likely resist curvature stress and impede the ion and water penetration required for structural rearrangement, rendering high-order particles refractory to release. This interpretation is consistent with reports linking less-ordered and hexagonal-phase LNP structures to improved knockdown efficiency^21,22^, and with analogous principles in viral uncoating, where productive delivery requires a finely tuned balance between extracellular stability and susceptibility to triggered disassembly^47^. Notably, classical rupture markers such as galectin recruitment do not reliably predict productive cytosolic release, even when endosomes remain perforated^13,48^. In this view, low-order packing represents an energetically primed configuration: stable enough to preserve particle integrity during trafficking yet sufficiently disordered to respond to endosomal acidification and enable efficient cytosolic access.

## Conclusion

The demonstration that mesoscale siRNA packing order, not lipid composition alone, is a primary determinant of LNP potency reframes a long-standing design challenge. Rather than screening lipid compositional space empirically, formulations can be rationally tuned to target internal architecture directly. Packing order thus is not just an incidental byproduct of formulation but can be utilized as a programmable property governed by ionizable lipid identity and electrostatic stoichiometry to optimize functional nucleic acid delivery. Our single-particle framework resolves this property at the individual-particle level, transforming it from an ensemble observation into a quantitative, controllable parameter, informing formulation design The actionable principle is clear: effective formulations should favor less compact, low-order assemblies that retain encapsulation efficiency while remaining structurally responsive to endosomal acidification, with composition and N/P ratio providing two independent handles for achieving this balance. Whether these structure-function relationships extend to alternative cargoes and in vivo contexts remains to be established, but the platform introduced here provides the quantitative foundation needed to answer such questions systematically.

## Supporting information

suppementary infromattion

## Acknowledgements

Work was funded by the Novo Nordisk foundation challenge center for Optimised Oligo escape (NNF23OC0081287), the center for 4D cellular dynamics (NNF22OC0075851) VILLUM FONDEN (40516 and 40801), and Lundbeck foundation Nanopans Project (R453-2024-359). N.S.H. is member of the Integrative Structural Biology Cluster at the University of Copenhagen.

## Author Contribution

AB, GK and NSH wrote the paper with feedback from all authors. GK and AB performed LNP formulation and characterization. GK performed spinning disk cell imaging with assistance from GB and FHS performed Lattice light sheet cell imaging with the help of AB and GK. GK, and FHS, handled cell culturing and plating with the help of GB and VP. GB generated HEK293T stably expressing d2eGFP with the help of VP. GK performed the cryo-EM experiments with the help of AB. AB developed the analysis algorithms and pipelines with help from GK. AB analyzed the data with the help of GK. MD prepared the lipid packing model and developed the analysis code with inputs from AB and GK. SM and NS performed preliminary experiments and analysis. NSH conceived the project, acquired funding and had the overall supervision and project management.

## Disclaimer

N.S.H. is the CSO and co-founder of EDGE Biotechnologies. A.B. is part time employee of EDGE Biotechnologies.

## Methods

### Materials

1,2-distearoyl-sn-glycero-3-phosphocholine (DSPC, 850365P), 1,2-dioleoyl-sn-glycero-3-phosphoethanolamine-N-[methoxy(polyethylene glycol)-2000] (ammonium salt) (PEG2000-PE, 880130P), and 1,2-distearoyl-sn-glycero-3-phosphoethanolamine-N-[biotinyl(polyethylene glycol)-2000] (ammonium salt) (PEG2000-DSPE-biotin, 880129P) were purchased from Avanti Polar Lipids (Alabaster, AL, USA). Cholesterol (C8667) was obtained from Sigma-Aldrich (St. Louis, MO, USA). 9-Heptadecanyl 8-{(2-hydroxyethyl)[6-oxo-6-(undecyloxy)hexyl]amino}octanoate (SM-102, HY-134541), ([(4-hydroxybutyl)azanediyl]di(hexane-6,1-diyl) bis(2-hexyldecanoate)) (ALC-0315, HY-138170), and 2-[(polyethylene glycol)-2000]-N,N-ditetradecylacetamide (ALC-0159, HY-138300) were obtained from MedChemExpress LLC (Monmouth Junction, NJ, USA). DPPE-ATTO488 (AD 488-151) and DPPE-ATTO655 (AD 655-151) were acquired from ATTO-TEC GmbH (Siegen, Germany). eGFP-targeting siRNA with Alexa Fluor 647 conjugated at the 3′ end of the sense strand (eGFPsiRNA_sense3′Alexa647N) was purchased from Integrated DNA Technologies (IDT, Coralville, IA, USA). Silencer™ eGFP siRNA (AM4626) was obtained from Invitrogen (Thermo Fisher Scientific, Waltham, MA, USA).

### LNP formulation and characterization

#### Microfluidics: LNP formulation

Ethanol solutions of the lipid mixes at indicated molar ration and total lipid concentration were prepared, as described in Supplementary Tables S1-3. The RNA was diluted in 10mM citrate buffer pH 4 (132-04-3, Thermo Fisher Scientific, Waltham, MA, USA). LNPs were prepared using a staggered herringbone micromixer (SHM) glass chip, with channel depth of 0.08mm (LTF-012.00-4264, Darwing Microfluidis, Paris, France) as the microfluidic device. The lipid mixture and the RNA solution were injected into the SHM via 250 μL PTFE Peek Syringes (10103660, Hamilton, Reno, NV, USA) and mixed at a N/P ratio of 3.

The SHM features three converging inlets, where the organic phase containing the lipid mixture flows through the two outer channels and the central channel delivers the aqueous phase comprising RNA. Flow rates were control by a Pump 33 dual drive system (703333, Harvard Apparatus, Massachusetts, USA). LNPs were collected at the outlet in 1.5 mL of 1X PBS PH 7.4 (AM9625, Invitrogen, Thermo Fisher Scientific, Waltham, MA, USA), to bring ethanol concentration below 5%. Residual ethanol was removed by overnight dialysis at 4 °C in 1X PBS using 100kDa molecular weight cut-off (MWCO) dialysis tubing (131420, Biotech, Repligen, USA). The LNP suspensions were concentrated by centrifugation at 4000 × g for 30 min at 4 °C using 10 kDa MWCO Amicon Ultra-4 centrifugal filter units (UFC8010, Merck Millipore, Burlington, MA, USA).

Encapsulated RNA was quantified using a modified Quant-iT™ RiboGreen™ RNA Assay (R11490, Invitrogen, Thermo Fisher Scientific, Waltham, MA, USA). A Triton-based calibration curve (1× TE + 0.5% Triton X-100) was generated using the same siRNA as in the LNP formulations (Silencer™ eGFP siRNA). Each sample was measured in parallel as free RNA (1× TE) and total RNA (1× TE + 0.5% Triton X-100). After incubation with RiboGreen working reagent, fluorescence was blank-corrected and converted to RNA concentration. Encapsulated RNA was determined as total minus free RNA, corresponding to the RNA protected within intact LNPs.

#### Dynamic light scattering: Size characterization of LNPs

The hydrodynamic diameter of lipid nanoparticles (LNPs) was measured by dynamic light scattering (DLS) using a Zetasizer Pro system (Malvern Panalytical, UK). Samples were diluted in 1X PBS to achieve comparable particle concentrations across formulations. For each LNP formulation, three independent technical replicates were prepared. Each replicate was measured three times consecutively. DLS data were analyzed as number-weighted size distributions. For each technical replicate, repeated runs were combined by calculating the mean number-percentage distribution (with SD) at each size bin, and this averaged distribution was fitted with a lognormal model. The distribution mode (reported particle size) was calculated from fitted parameters as *exp(μ−σ*^*2*^). Final values are reported as mean ± SD across technical replicates.

#### Cryogenic Transmission Electron Microscopy

Lipid nanoparticles (LNPs) were concentrated to a final total lipid concentration of ~15 mg/mL. Lacey carbon 300 mesh copper grid (Ted Pella Inc., Redding, CA, USA) was glow discharged on Ace 200 Leica glow discharge unite (30 sec at 15mA). A 3μL aliquot of the suspension was applied to the grid and plunge-frozen using a Mark IV Vitrobot (FEI, Hillsboro, OR), with a blot-force of 4 for 3 seconds to generate vitreous ice. Grids were stored in liquid nitrogen prior to imaging. Cryogenic transmission electron microscopy (cryo-TEM) was performed using a Tecnai G2 20 microscope (Thermo Fisher Scientific, USA) operated at 200 kV under low-dose rate. Images were acquired with a bottom mounted FEI High-Sensitive (HS) 4k × 4k Eagle camera. All sample preparation and imaging were carried out at the Core Facility for Integrated Microscopy, University of Copenhagen (Copenhagen, Denmark).

#### Total Internal Reflection Fluorescence microscopy: Single-particle fluorescence assays

Single-particle experiments were conducted using an inverted total internal reflection fluorescence (TIRF) microscope, model IX83 (Olympus). The system was equipped with an electron-multiplying charge-coupled device (EMCCD) camera (ImagEM X2, Hamamatsu), a 100× oil immersion objective lens (UAPON 100XOTIRF, NA 1.49, Olympus) and a quad-band emission filter cube designed to remove excitation laser light from the detection pathway. Fluorescence excitation was achieved using two solid-state lasers (Olympus) operating at wavelengths of 488 nm (Atto488-labeled lipids) and 640 nm (Alexa647-labeled cargo). Prior to acquisition, optimal focus and TIRF angle were established. Data were acquired with a 200nm evanescent field penetration depth, an EM gain setting of 300, and image resolution of 512 × 512 pixels in 16-bit grayscale. Each acquired field of view corresponds to a physical area of 81.92 μm^2^.

#### TIRF: High-throughput single-particle characterization of LNPs

Plasma activated glass slides (80608, Ibidi, Martinsried, Germany) incorporating sticky-Slide VI 0.4 (10812, Ibidi, Martinsried, Germany) were functionalized using PLL-g-PEG and PLL-g-PEG-biotin (Susos AG, Dübendorf, Switzerland) at a 100:1 ratio. A neutravidin (11544037, Fisher Scientific, Waltham, MA, USA) layer was subsequently applied to enable specific immobilization of biotinylated particles.

Biotinylated LNPs diluted in 1X PBS buffer pH 7.4 were injected into the microfluidic channel and incubated for 10 min to allow surface immobilization. The concentration was adjusted to achieve densities of approximately 40-70 LNPs per field of view. Unbound particles were removed by rinsing the channels with 1X PBS (5 × 100 μL).

Each high-throughput assay, was performed in four independent technical replicates, comprised imaging of ~14,400 individual fields of view, corresponding to approximately 0.5-1 million single particles per LNP formulation.

#### TIRF: High-throughput imaging of single-molecule RNA

Plasma activated glass slides were assembled with sticky-Slide VI 0.4 microfluidic chambers. Fluorescently labeled RNA, diluted in 1X PBS (pH 7.4), was introduced into the channels and incubated for 10 min at room temperature to allow surface immobilization. The RNA concentration was optimized to achieve a low surface density, so as to attain spatially distinct single-molecule. Unbound molecules were removed by washing the channel with 1X PBS (10 × 100 μL). Each experiment was performed in four independent replicates, comprising imaging of ~1200 distinct fields of view.

### Live-cell assays

#### Generation of d2eGFP-expressing cells

HEK293 cells stably expressing destabilized enhanced green fluorescent protein (HEK293-d2eGFP) were established by lentiviral transduction with a vector encoding d2eGFP. Lentiviral particles were generated by co-transfecting HEK293T packaging cells with pMD2.G, psPAX2, and pLV-CMV-d2eGFP plasmids using Lipofectamine 3000 (Thermo Fisher Scientific, Waltham, MA, USA) according to the manufacturer’s protocol.

#### Cell culture

HEK293-d2eGFP cells were cultured in Nunc EasYFlask 25 cm^2^ flasks in Dulbecco’s Modified Eagle’s Medium (DMEM, 4500 mg/L glucose) supplemented with 2 mM L-glutamine, 1 mM sodium pyruvate, and 10% (v/v) heat-inactivated fetal bovine serum (FBS). Cells were incubated at 37 °C in a humidified incubator with 5% CO2 and 70% relative humidity and passaged every 3-5 days. All experiments were performed using cells between passages 15 and 20.

#### Spinning Disk Confocal (SDC) Microscopy: Live-cell d2eGFP knockdown assay

Live-cell imaging was performed on a spinning disk confocal microscope equipped with a 20× air objective lens (Olympus, NA = 0.5), a Prime 95B CMOS camera (Teledyne Photometrics, effective pixel size of 183 nm × 183 nm) and an environmental control chamber (37 °C, 5% CO2, humidity).

For knockdown assays, cells were seeded at a density of 10,000 cells per well in Ibidi 8-well μ-slides (Ibidi, Martinsried, Germany) containing 200 μL of complete DMEM and incubated for 48 h prior to treatment. Prior to imaging, cells were incubated with 200 μL of imaging medium (phenol red-free DMEM supplemented with 10% FBS) containing LNPs (Supplementary Table S3) encapsulating siRNA targeting eGFP (final concentration 300 nM of encapsulated siRNA), or with imaging medium alone as a negative control. Imaging was initiated 1 h 15 min after LNP addition and continued every 20 min for 24 h. For each condition, 10 positions per well were imaged in parallel. At each position, seven z-plane images were collected with 3 μm spacing using a 488 nm excitation laser at 20% laser power with an exposure time of 50.04 ms. All acquisition settings were kept constant across experiments. Each condition was performed in three independent technical replicates.

#### Lattice Light-Sheet (LLS) Microscopy

Experiments were performed on a ZEISS Lattice Lightsheet 7 microscope equipped with a 13.3× illumination objective (NA 0.4), a 44.8× detection objective (NA 1.0), and two Hamamatsu ORCA-Fusion sCMOS cameras (6.5 μm pixel size). The light sheet was configured with a Sinc^3^ profile of 30 × 1000 μm and cropped to 296 μm in width and 43.5 μm in height at a 30° angle, corresponding to 37.1 μm perpendicular to the sample surface. Z-stacks were acquired with a step size of 200 nm. PSF calibration in the Z-direction was performed using fluorescent LNPs in agarose, ensuring a full-width half maximum (FWHM) below 1.0 μm for the 640 nm excitation laser.

#### LLS: PSF calibration and single-particle characterization

For PSF calibration and single-particle characterization, LNPs were suspended in 1.5% high-melt agarose prepared at 90 °C. A 1:50 dilution of LNP stock was mixed with molten agarose and cast into wells of glass-bottom Ibidi 8-well μ-slides. After cooling to form gels, samples were imaged under the same acquisition settings as live-cell experiments. Immobilized LNPs were excited with a 640 nm laser at either 30% or 100% power (10 ms exposure). Each condition was recorded in technical triplicates, with each replicate comprising 56 fields of view spanning the full sample well.

#### LLS: Live-cell internalization assay

For lattice light-sheet microscopy (LLSM) internalization assay, 10,000 cells were seeded into each well of fibronectin-coated (11 μg/mL) glass-bottom Ibidi 8-well μ-slides (Ibidi, Martinsried, Germany) and incubated for 24 h prior to treatment.

Prior to imaging, cells were treated with imaging medium (IMEM) containing either Comp. A N/P 3 LNPs, Comp. B N/P 3 LNPs (Supplementary Table S3), or no LNPs as control for 150 minutes. Following incubation, cells were washed twice with 200 μL imaging medium to remove unbound LNPs and maintained in imaging medium for live-cell imaging.

Live cell imaging was performed at 37°C under 5% CO2, and 70% humidity. LNPs were imaged using a 640 nm laser at 30% or 100% power with a 10 ms exposure time, while cytosolic d2eGFP was excited using a 488 nm laser at 4% power with an exposure time of 10 ms. Each condition was recorded in three independent biological replicates, with 12 fields of view per replicate.

### Data analysis and computational experiments

#### TIRF microscopy data processing and quantitative analysis

Quantitative analysis of single-particle TIRF datasets was performed through a multi-stage image processing pipeline designed to extract physical (Diameters) and molecular parameters (Copies of siRNA) of individual lipid nanoparticles. The workflow included correction for non-uniform illumination, detection and fitting of individual point spread functions (PSFs) to obtain integrated fluorescence intensities, pairwise colocalization of membrane and cargo signals to identify loaded particles, and quantitative conversion of fluorescence values into particle size and siRNA copy number. These steps enabled direct mapping of LNP size to cargo content at single-particle resolution.

#### Illumination correction

To compensate for spatial variations in excitation intensity across the TIRF field, raw fluorescence movies were corrected using a custom flat-fielding procedure. For each dataset, the mean projection along the temporal axis was smoothed with a Gaussian filter (σ ≈ 30 pixels) and fitted with a two-dimensional Gaussian function combined with a sloped planar background. This composite model captures both the radially decaying excitation characteristic of objective-based TIRF illumination and small linear gradients introduced by slight beam misalignment or optical throughput variations. Each movie frame was divided by the resulting normalized illumination profile, and the corrected stack was rescaled to preserve the original mean intensity.

Without correction, PSFs located near the edges of the field experience systematically lower apparent brightness compared to those near the center, leading to an artificial broadening of intensity distributions and bias in quantitative single-particle analyses. Applying the illumination correction homogenized the apparent intensity of PSFs across the field of view, enabling accurate comparison of integrated fluorescence values between particles.

#### Single-particle detection and Gaussian fitting of PSFs

Individual particles were identified in each frame using a GPU-accelerated Laplacian-of-Gaussian–based detector optimized for diffraction-limited features. For every detected particle, a 7 × 7 pixel region (corresponding to 1.12 μm × 1.12 μm at 160 nm px^−1^) was fitted with a two-dimensional Gaussian function plus a constant background using nonlinear least-squares optimization. This window fully encompasses the expected point-spread function under the imaging conditions.

The fitted parameters included the particle center coordinates (*x*_0_, *y*_0_), the Gaussian widths (*σ*_*x*_, *σ*_*y*_), and the amplitude *A*. From these, the integrated intensity of each particle was calculated as

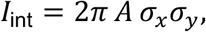

which was used as the sole quantitative descriptor of particle brightness in subsequent analyses. To avoid artifacts from detector saturation, oversaturation masks generated from the raw movies were used to flag particles located within 3 pixels of saturated regions. All results were saved separately for the membrane and cargo channels and used for downstream intensity-based comparisons between individual LNPs.

#### Colocalization of membrane and cargo particles

To quantify the fraction of LNPs containing encapsulated cargo, colocalization analysis was performed between particles detected in the membrane and cargo fluorescence channels. Each field of view was analyzed pairwise, matching particles based on spatial proximity. Membrane and cargo particle centroids were considered colocalized when their lateral distance was below 1 pixel (160 nm), corresponding to a conservative estimate of the typical localization accuracy expected under the imaging conditions to minimize false positives.

For each LNP formulation and replicate, the total number of membrane particles and the number of colocalized membrane-cargo pairs were quantified. The colocalization fraction was calculated as the ratio of colocalized particles to the total number of membrane-associated particles. Integrated intensities from the fitted Gaussians were then used to compare the fluorescence distributions of colocalized and non-colocalized populations.

#### Quantification of siRNA copies per LNP

To quantify siRNA loading per particle, fluorescent intensities were transformed into physical LNP radii and discrete siRNA copy numbers. Colocalization of siRNA with LNPs was first used to identify cargo-bearing particles. Subsequent analysis, apart from the intensity to size conversion in LNPs, was restricted to this subset.

Integrated LNP fluorescence intensity, I, was converted to particle radius, r, using a dynamic light scattering (DLS) anchor. Assuming fluorescence scales with surface area,

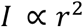

the proportionality constant

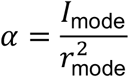

was defined from the modal intensity of the LNP distribution and the corresponding DLS-derived hydrodynamic radius. Each LNP was then assigned an estimated size:

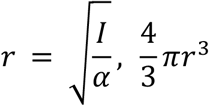

Cargo channel intensities were converted to molecule counts using single-molecule siRNA calibration data, where a gaussian mixture model in log space identified the monomer peak, yielding mean single-molecule intensity *I*_1_. For each particle, siRNA copy number was estimated as

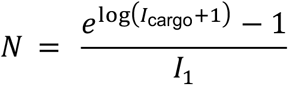

This procedure produces a direct mapping of siRNA copy number as a function of LNP radius, enabling quantitative analysis of size-dependent loading.

#### Extracting multimodal scaling laws

With siRNA counts mapped to LNP radius, 2D histograms revealed distinct ridges of density following different scaling exponents. We first attempted fitting parameterized multi-component distributions directly in the 2D plane, but this approach was unstable. Instead, we adopted a band-wise analysis: LNPs were stratified into narrow radius intervals, and siRNA copy-number distributions were examined within each band, as depicted in Fig. 3.

Across conditions, siRNA counts within size bands were frequently bimodal. To assess this formally, we fitted one-vs.\ two-component gaussian models in log space and compared them using the Bayesian Information Criterion (BIC). Approximately half of the conditions generally favored size-bands with a two-component fit; the remainder were best described by a single gaussian.

Scaling laws were therefore evaluated in two complementary ways:

a. the mean siRNA count per LNP radius (robust to uni- or bimodality), and
b. separate scaling exponents for “high-load” and “low-load” subpopulations, by tracking which gaussian corresponded to the higher vs. lower siRNA peak within each radius band.

#### Simulating scaling laws via stochastic LNP mergers

To interpret observed scaling of siRNA loading as a function of LNP size, we implemented a stochastic coagulation model representing the dynamic formation of LNPs through successive binary mergers. Each event corresponds to the fusion of two lipid domains, consistent with proposed LNP self-assembly mechanisms based on progressive domain coalescence^49,50^. The process is simulated as an asynchronous, event-driven system following Gillespie dynamics, ensuring a realistic evolution of the particle size distribution over time.

At the initial time point (*t* = 0), *N*_0_ spherical LNPs are generated with radii drawn from a normal distribution (*r*_0_ ± *σ*_*r*_). Each particle is assigned a discrete siRNA cargo, initialized according to a Bernoulli distribution that determines the encapsulation fraction:

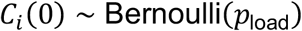

where *p*_load_ defines the initial probability that an LNP contains siRNA (i.e., 1 − *p*_load_ is the empty fraction).

At each stochastic event, two particles, i and j, are selected uniformly at random and merged into a new particle. The resulting LNP inherits its physical and cargo properties through additive combination: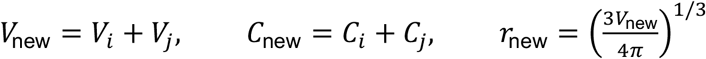

The merged particle replaces the two parent particles in the active population, and the process continues until the largest particle reaches a predefined hydrodynamic radius (*r*_max_ = 1500nm), corresponding to the upper experimental size regime. Volume-additivity was selected as electron-microscopy indicated that the lipids aggregate in a manner consistent with volumetric coordination.

The remaining particles at this stopping point represent the terminal distribution of LNP sizes and siRNA copy numbers, *p*_*τ*_ (*r, C*), at time *τ*. This ensemble provides a synthetic counterpart to experimentally measured populations, where individual nanoparticles are observed after assembly rather than during formation.

From this terminal distribution, the empirical relationship between siRNA count and LNP radius is determined by fitting a power law of the form

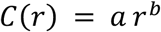

where *b* denotes the effective scaling exponent. For homogeneous seeding and conserved cargo, *b* approaches 3, reflecting volume-proportional loading. Under sparse encapsulation (*p*_load_ ≪ 1), the effective exponent decreases, typically near *b* ≈ 1.2 to *b* ≈ 2.1 for *p*_load_ = 0.05 to *p*_load_ = 0.2, respectively, consistent with experimental measurements. This deviation from direct volumetric scaling arises naturally from stochastic variations in merger history and the low frequency of cargo-bearing progenitors, without requiring any explicit cargo loss or surface-limited loading assumptions. Our computational coalescence modeling as such aims to interpret where the observed scaling exponents arise from. While a uniform kernel, that is the probability of any two particles merging is the same for any other, does not explain the observed heterogeneity in scaling for some conditions, we are able to theorize that the sub-volume proportional cargo loading may very well arise from the initial encapsulation efficiency.

#### Live-cell quantification of siRNA-mediated eGFP knockdown following LNP delivery

Knockdown kinetics were quantified from time-lapse fluorescence measurements acquired by spinning-disk confocal microscopy. For each well and imaging position, seven z-planes (3 μm spacing) were acquired and collapsed by maximum-intensity projection. Cells were segmented on maximum-intensity projections of the eGFP channel together with a user-selected brightfield z-plane to improve boundary detection when fluorescence was low after knockdown.

Within each field of view (FOV) and frame, mean cellular eGFP intensity was computed per segmented cell; these per-cell values were averaged to yield an FOV-level mean for that frame. For a given replicate and frame, the replicate-level mean was then obtained by averaging the FOV-level means across all FOVs imaged for that replicate. This produced one time series per replicate, *T*_*i*_ (*t*), sampled every 20 min for ~24 h after LNP treatment.

To construct a global reference trajectory for untreated controls, all control replicates from the same experiment were pooled to form *C*(*t*), the inverse-variance–weighted mean of per-replicate control curves:

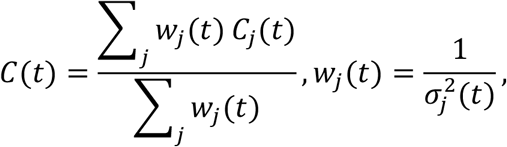

where *C*_*j*_ (*t*) and *σ*_*j*_ (*t*) denote the mean and standard deviation of the control replicate *j* at time *t*. The pooled control trajectory captures the shared recovery and bleaching dynamics of untreated cells.

Each treated replicate *T*_*i*_ (*t*) was normalized to the control by first estimating a scalar baseline-alignment factor *s*_*i*_ that minimized the least-squares difference to *C*(*t*) over the initial “no-effect” window (within the first hour):

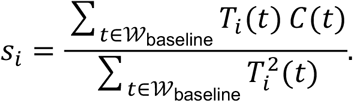

The aligned trace was

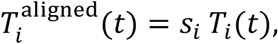

and the normalized signal ratio

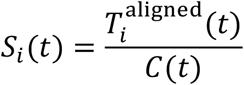

was computed at all shared time points. By construction, *S*_*i*_ (*t*) ≈ 1 at early times and decreases when eGFP expression is suppressed relative to control.

To suppress frame-to-frame noise and enforce the expected monotonic decline of fluorescence as siRNA knockdown progresses, *S*_*i*_ (*t*) was first smoothed with a moving-average filter and then projected onto a monotone non-increasing function using isotonic regression. When available, propagated per-timepoint uncertainties were used as weights in the fit. The resulting piecewise-constant curve 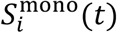 represents the denoised knockdown trajectory for that replicate.

From 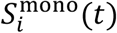, the maximum knockdown was calculated as:

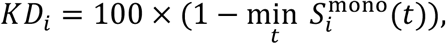

reflecting the largest fractional decrease in eGFP intensity relative to control. All analyses were performed independently per replicate (one well). Replicate-level values were summarized per LNP formulation as mean ± SEM.

### Quantification of internalized LNPs per cell

To quantify the average number of internalized LNPs per cell, 3D fluorescence volumes acquired in the lattice light-sheet microscopy (LLSM) assay were analyzed across all biological replicates. Endosomes can undergo rapid fusion and fission that can result in in multiple LNPs per endosome. This consequently challenges quantitative extraction of the LNP number per cell. To extract therefore an estimate of the number of LNPs internalized per cell we quantified the total background-corrected cellular fluorescence of LNPs in cells and divided with the corresponding mean single-LNP intensity.

Cells expressed cytosolic d2eGFP, which was used for segmentation of individual cellular volumes. A representative z-slice containing well-separated cells was segmented using Cellpose-SAM, and segmentation was propagated across the remaining z-slices using SAM2 by interpreting each slice as a temporal variation of the reference plane. The resulting masks were combined to generate a 3D cell volume for each cell.

For each cell, the mean per-voxel fluorescence intensity of the LNP channel was measured within the segmented volume. These per-cell values were then averaged across all cells within each biological replicate to obtain the replicate-level mean per-voxel fluorescence intensity. The same analysis was applied to control samples (no-LNP) to determine cellular background fluorescence. The background-corrected LNP-associated intensity per voxel was then obtained by subtracting, for each replicate, the mean per-voxel intensity measured in the control condition from that measured in the corresponding experimental condition. The resulting background-corrected intensity was summed across all voxels within each cell, yielding the total fluorescence associated with internalized LNPs.

Single-particle calibration was performed using immobilized LNPs embedded in 1.5% high-melt agarose and imaged under identical acquisition conditions. Particles were detected using a Crocker-Grier-based 3D detection algorithm, and each detection was fitted with a 3D Gaussian model to obtain the integrated fluorescence intensity. The mean integrated intensity of isolated LNPs was used as the reference brightness of a single particle for each formulation.

For each experimental condition, the total background-corrected cellular fluorescence was divided by the corresponding mean single-LNP intensity, providing an estimate of the number of LNPs internalized per cell. The resulting values were averaged across all cells within each biological replicate, and the final mean and standard error of the mean (SEM) were computed across replicates.

## Notes

### Summary of Updates

minor modification in one figure and text editing/sharpening

