## Supplementary material for "Tuning siRNA packing order in lipid nanoparticles modulates oligonucleotide functional delivery": suppementary infromattion

**Supplementary Table S1: Formulation and imaging parameters for size-calibrated fluorescent LNPs.** All formulations share Composition A (SM102/DSPC/cholesterol/ $PEG_{2000}$-PE/$PEG_{2000}$-DSPE-biotin/DPPE-ATTO488, 50:9.5:38.5:1:0.5:0.5 mol%) and encapsulate eGFP siRNA (sense-3′-Alexa 647N) at N/P = 3. Particle size was tuned by varying the microfluidic parameters and lipid concentration listed below. DLS size values are reported as mean ± s.d. (n = 3 independent preparations). DLS, dynamic light scattering; TFR, total flow rate; FRR, flow rate ratio; TIRF, total internal reflection fluorescence.

| | **LNP 1** | **LNP 2** | **LNP 3** | **LNP 4** | **LNP 5** | **LNP 6** | **LNP 7** |
|---|---|---|---|---|---|---|---|
| **DLS size (nm)** | 39 ± 2 | 54 ± 2 | 64 ± 3 | 76 ± 5 | 78 ± 2 | 99 ± 3 | 102 ± 2 |
| ***Microfluidics*** | | | | | | | |
| Lipid conc. (mg $mL^{-1}$) | 1 | 1 | 5 | 7 | 10 | 7 | 14 |
| TFR (µL $min^{-1}$) | 666 | 100 | 100 | 100 | 100 | 100 | 100 |
| FRR | 3 | 3 | 3 | 3 | 3 | 1 | 1 |
| ***TIRF parameters*** | | | | | | | |
| **Laser emission** | *Laser power (%) / exposure time (ms)* | | | | | | |
| $\lambda_{ex}$ = 488 nm | 8.5 / 1,000 | 8.5 / 750 | 8.5 / 300 | 8.5 / 300 | 8.5 / 250 | 8.5 / 200 | 8.5 / 100 |
| $\lambda_{ex}$ = 640 nm | 5 / 1,000 | 5 / 750 | 5 / 300 | 5 / 300 | 5 / 200 | 5 / 150 | 5 / 50 |

**Supplementary Table S2: LNP formulations used to assess subpopulation coexistence across compositions and N/P ratios.** Compositions A and B differ primary in the identity of the ionizable lipid (SM-102 and ALC-0315, respectively), with a minor shift in the ionizable-lipid-to-cholesterol molar ratio while all remaining components are matched. All formulations were prepared under identical microfluidic conditions (total lipid concentration 5 mg $mL^{-1}$, TFR 100, FRR 3). TIRF parameters were optimized per formulation to ensure comparable signal-to-noise conditions. DLS size values are reported as mean ± s.d. (n = 3).

| | **Comp. A NP1** | **Comp. A NP3** | **Comp. A NP6** | **Comp. B NP3** |
|---|---|---|---|---|
| **DLS size (nm)** | 74 ± 1 | 71 ± 2 | 70 ± 5 | 65 ± 3 |
| ***Composition (mol%)*** | | | | |
| SM-102 | 50 | 50 | 50 | – |
| ALC-0315 | – | – | – | 46.3 |
| DSPC | 9.5 | 9.5 | 9.5 | 8.9 |
| Cholesterol | 38.5 | 38.5 | 38.5 | 42.7 |
| $PEG_{2000}$-PE | 1 | 1 | 1 | – |
| ALC-0159 | – | – | – | 1.1 |
| $PEG_{2000}$-DSPE-biotin | 0.5 | 0.5 | 0.5 | 0.5 |
| DPPE-ATTO488 | 0.5 | 0.5 | 0.5 | 0.5 |
| DPPE-ATTO655 | – | – | – | – |
| ***Cargo*** | | | | |
| eGFP siRNA | Sense-3′-Alexa 647N | Sense-3′-Alexa 647N | Sense-3′-Alexa 647N | Sense-3′-Alexa 647N |
| N/P ratio | 1 | 3 | 6 | 3 |
| ***TIRF parameters*** | | | | |
| **Laser emission** | *Laser power (%) / exposure time (ms)* | | | |
| $\lambda_{ex}$ = 488 nm | 10 / 200 | 10 / 100 | 10 / 75 | 10 / 40 |
| $\lambda_{ex}$ = 640 nm | 12 / 12 | 12 / 12 | 12 / 12 | 12 / 10 |

**Supplementary Table S3: Composition and N/P ratio effect in live cell assays.** Compositions A and B differ primarily in the identity of the ionizable lipid (SM-102 and ALC-0315, respectively), with a minor shift in the ionizable-lipid-to-cholesterol molar ratio while all remaining components are matched. All formulations were prepared under identical microfluidic conditions (total lipid concentration 5 mg mL$^{-1}$, TFR 100, FRR 3). DLS size values are reported as mean ± s.d. (n = 3).

| | **Comp. A NP1** | **Comp. A NP3** | **Comp. A NP6** | **Comp. B NP3** |
|---|---|---|---|---|
| **DLS size (nm)** | 84 ± 2 | 77 ± 1 | 63 ± 1 | 78 ± 2 |
| ***Composition (mol%)*** | | | | |
| SM-102 | 50 | 50 | 50 | – |
| ALC-0315 | – | – | – | 46.3 |
| DSPC | 9.5 | 9.5 | 9.5 | 8.9 |
| Cholesterol | 38.5 | 38.5 | 38.5 | 42.7 |
| $PEG_{2000}$-PE | 1 | 1 | 1 | – |
| ALC-0159 | – | – | – | 1.1 |
| $PEG_{2000}$-DSPE-biotin | 0.5 | 0.5 | 0.5 | 0.5 |
| DPPE-ATTO488 | – | – | – | – |
| DPPE-ATTO655 | 0.5 | 0.5 | 0.5 | 0.5 |
| ***Cargo*** | | | | |
| Silencer™ eGFP siRNA | All formulations | | | |
| N/P ratio | 1 | 3 | 6 | 3 |

**Supplementary Note 1: Size-dependent siRNA partitioning between formulations.** Stratifying total siRNA content into five diameter intervals revealed a consistent depletion of Composition A relative to Composition B at smaller and mid-sized particles (0.42× at 0–75 nm; 0.58× at 75–100 nm; 0.72× at 100–180 nm; 0.63× at 180–320 nm), with inversion only in the largest bin (>320 nm; 3.15×). While bulk dosing based on input RNA concentration captures differences in mean loading, it fails to resolve this size-dependent cargo partitioning, which may have important implications for functional delivery, as discussed in the main text.

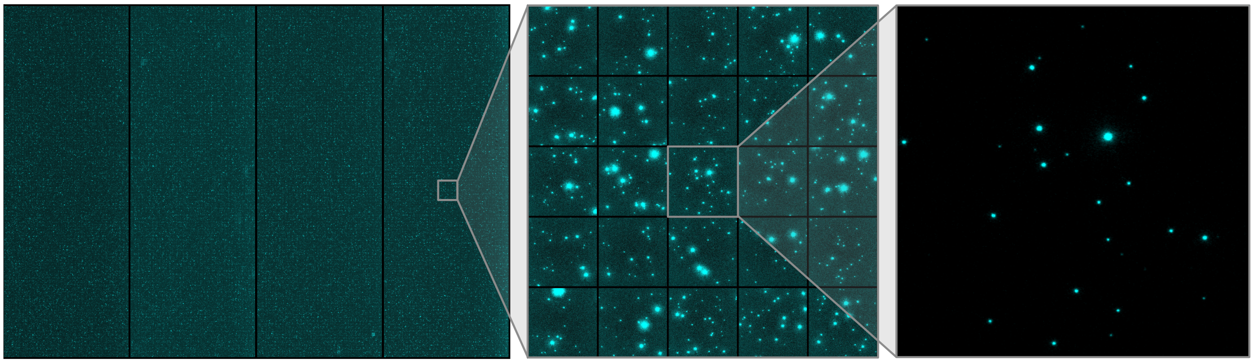

**Supplementary Figure S1 | High-throughput single-particle imaging of lipid nanoparticles.** Total internal reflection fluorescence (TIRF) montage of 4 replicates for a total of 14,400 fields of view (FOVs) acquired in the lipid fluorescence channel (ATTO488, cyan), representing more than 0.5 million individual LNPs. Progressive zoom-ins from the full dataset (left) through an intermediate grid of FOVs (middle) to a single representative FOV (right) illustrate the imaging scale. Dynamic contrast of the left and middle panels has been adjusted for visualization clarity. Each diffraction-limited spot corresponds to a single LNP. These data underpin the fluorescence–diameter calibration and single-particle heterogeneity analyses presented in the main text.

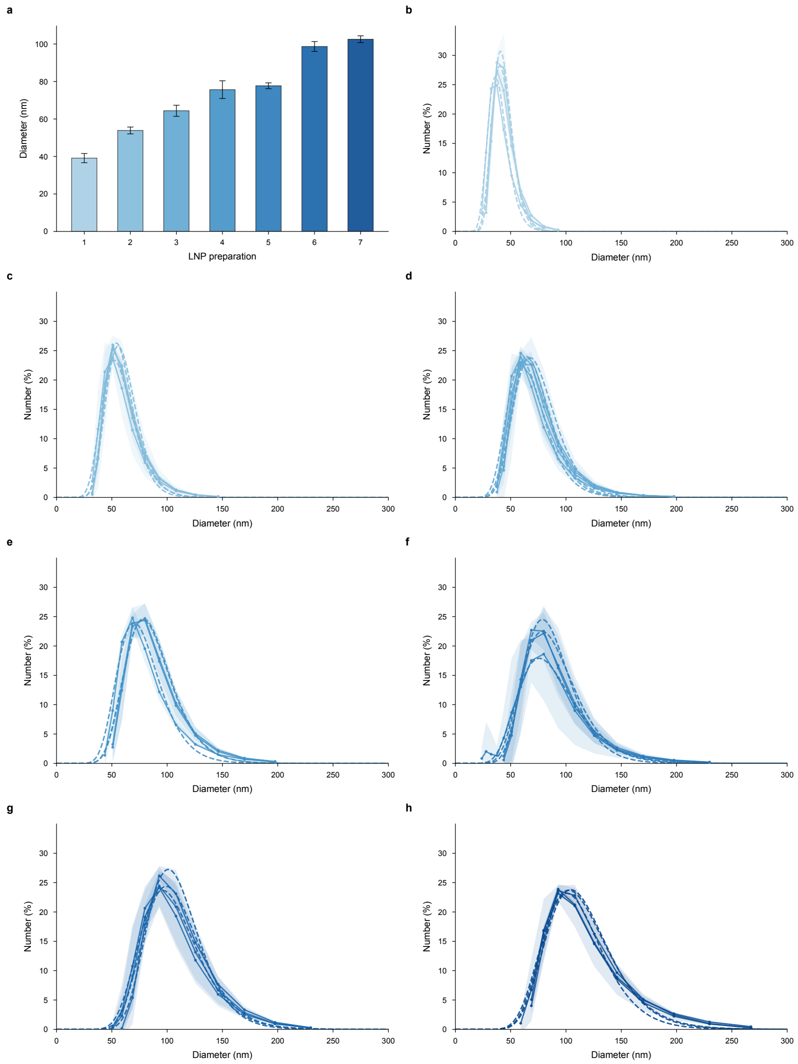


**Supplementary Figure S2 | Bulk size characterization of LNP formulations by dynamic light scattering. a**, Mode hydrodynamic diameters of the seven LNP preparations used for single-particle fluorescence calibration (Supplementary Table S1). Bars represent the mode diameter from number-weighted size distributions; error bars denote s.d. (*n* = 3–4 technical replicates). Bar shading progresses from light blue (preparation 1, smallest) to dark blue (preparation 7, largest). **b**–**h**, Number-weighted size distributions for preparations 1–7, respectively. Individual replicates are shown as dashed lines; the shaded envelope indicates ± 1 s.d. across replicates. Distributions are monomodal across all formulations, with mode diameters spanning approximately 40–110 nm. These bulk measurements provided the physical reference for converting fluorescence intensities into particle size.

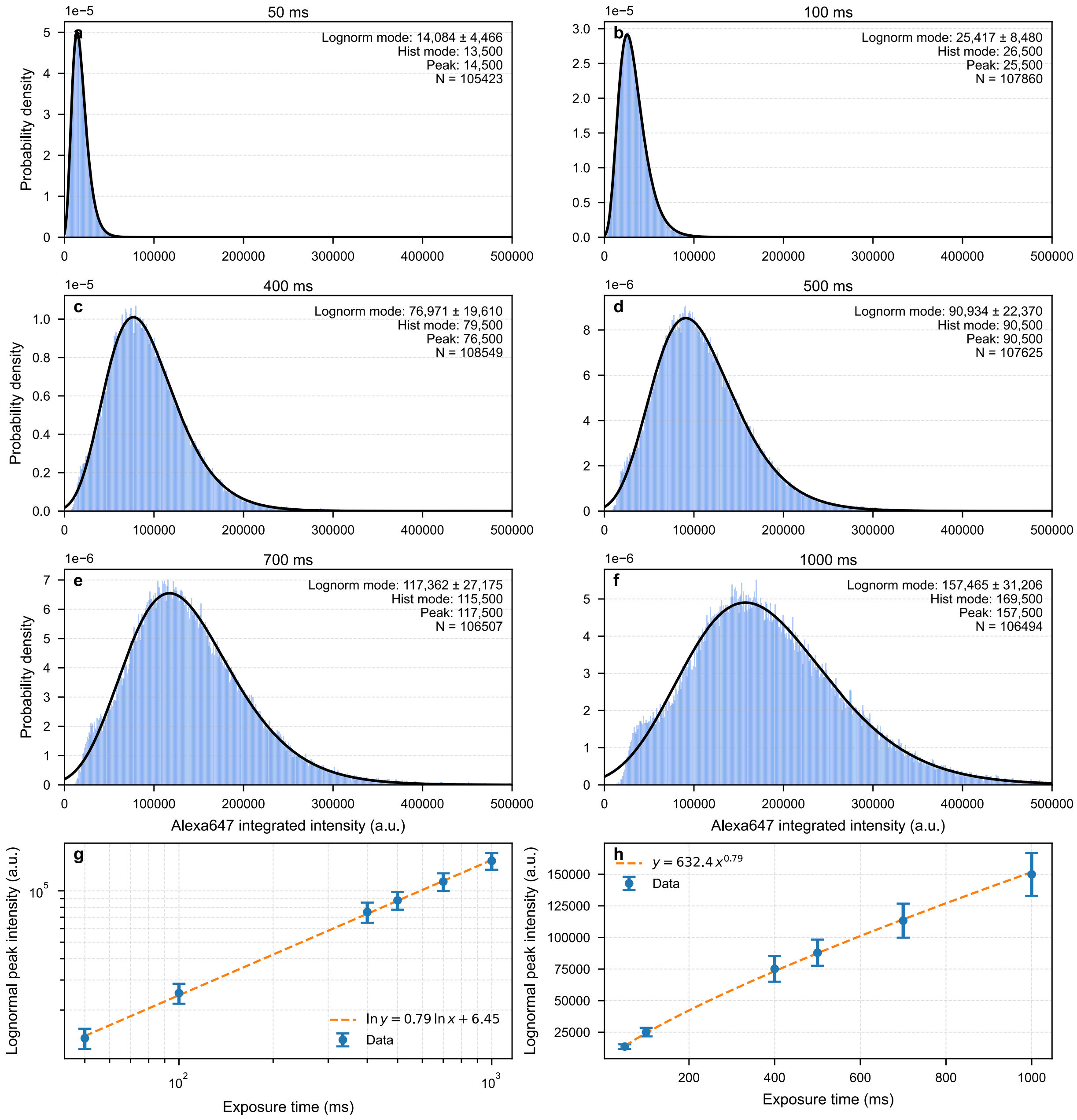


**Supplementary Figure S3: Exposure-time dependence of single-molecule Alexa 647 fluorescence intensity. a–f**, Single-molecule intensity distributions of Alexa 647-biotin immobilized on PLL-PEG-biotin/NeutrAvidin-functionalized coverslips, imaged under 640 nm excitation at six exposure times (50, 100, 400, 500, 700 and 1,000 ms). Light blue histograms show the integrated fluorescence intensity of individual molecules; black curves are log-normal fits. Insets report the log-normal mode ± s.d., histogram mode, peak value and number of molecules analysed (N > 105,000 per condition; three independent experiments). **g**, Log–log plot of log-normal mode intensity versus exposure time. Blue circles indicate the mode; error bars denote s.d. across three independent experiments. The orange dashed line is a linear fit in natural-log space (ln y = 0.79 ln x +

6.45). h, Same data displayed on linear axes with the corresponding power-law fit ($y = 632.4\ x^{0.79}$), showing a sublinear increase in detected signal with exposure time. This calibration was used to correct exposure-dependent intensity deviations in downstream siRNA quantification.

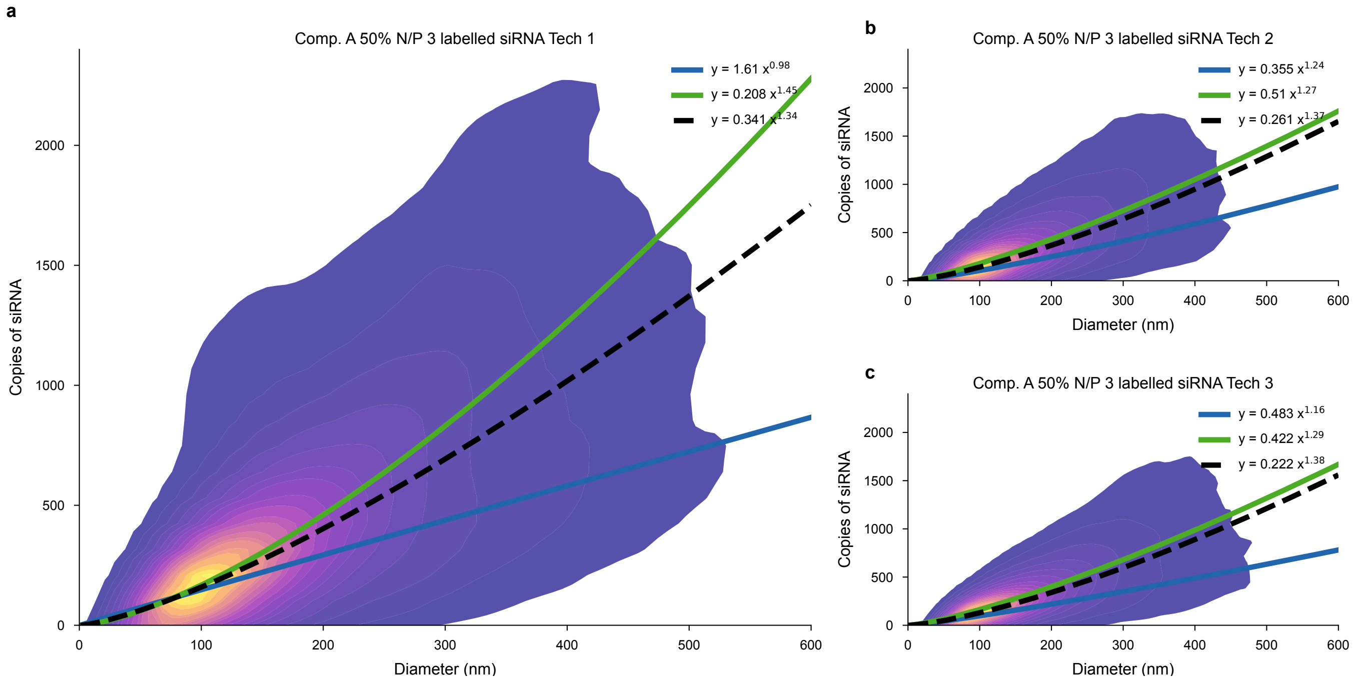


**Supplementary Figure S4: Control for fluorophore self-quenching in single-particle siRNA loading measurements. a–c**, Two-dimensional kernel density estimate (KDE) contour plots of labelled siRNA copy number versus LNP radius for Composition A (N/P 3) formulated with 50% Alexa 647N-labelled and 50% unlabelled eGFP-targeting siRNA, shown for three independent technical replicates (approximately 100,000–200,000 particles per replicate). Dashed lines represent power-law fits ($y = Ax^B$) to the low-order subpopulation (blue), the high-order subpopulation (green) and the overall population (black). The joint size-loading distributions consistently resolve two distinct packing subpopulations across all replicates, demonstrating that the observed bimodality does not arise from fluorophore self-quenching but reflects intrinsic heterogeneity in siRNA packing within the LNP population.

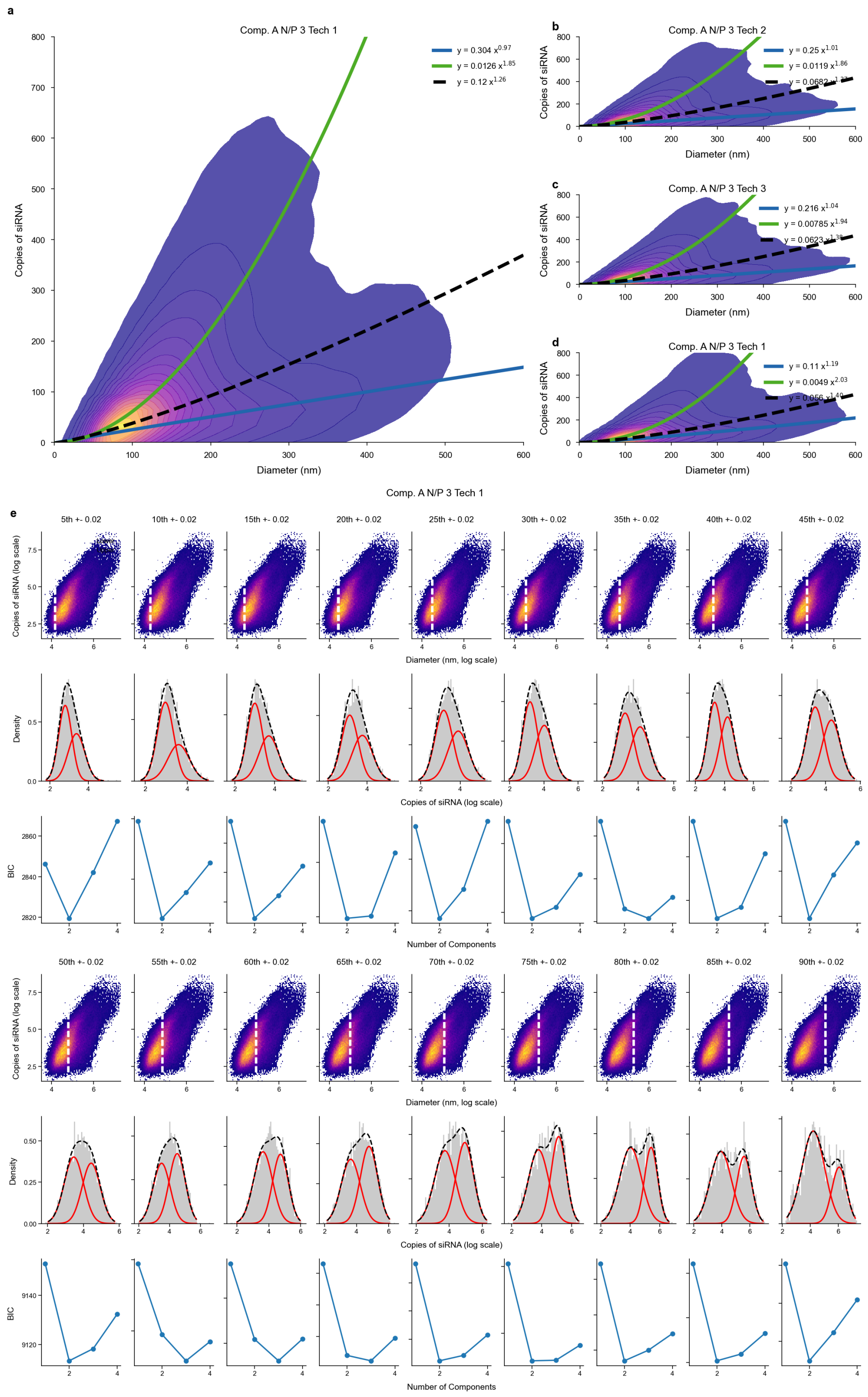


**Supplementary Figure S5: Replicate analysis of siRNA packing subpopulations for Composition A (N/P 3). a-d**, Two-dimensional kernel density estimates of siRNA copy number versus LNP diameter for four independent technical replicates (approximately 100,000–200,000 particles per replicate). Power-law fits ($y = Ax^B$) are shown for the low-order subpopulation (blue), the high-order subpopulation (green) and the overall population (black dashed). All replicates consistently resolve two distinct loading subpopulations. **e**, Gaussian mixture model (GMM) analysis of replicate 1, shown as representative, across 18 diameter percentile bins. Top row: Scatter plots of siRNA copy number versus diameter (log scale), colored by local density (purple to yellow); white dashed lines mark the percentile bin whose siRNA copy-number data are shown in the corresponding histogram below. Middle row: log-transformed siRNA copy-number distributions within each bin (grey histograms) overlaid with individual Gaussian components (red curves) and the summed fit (black curves). Bottom row: Bayesian information criterion (BIC) as a function of the number of mixture components (1–4); the minimum indicates the preferred model.

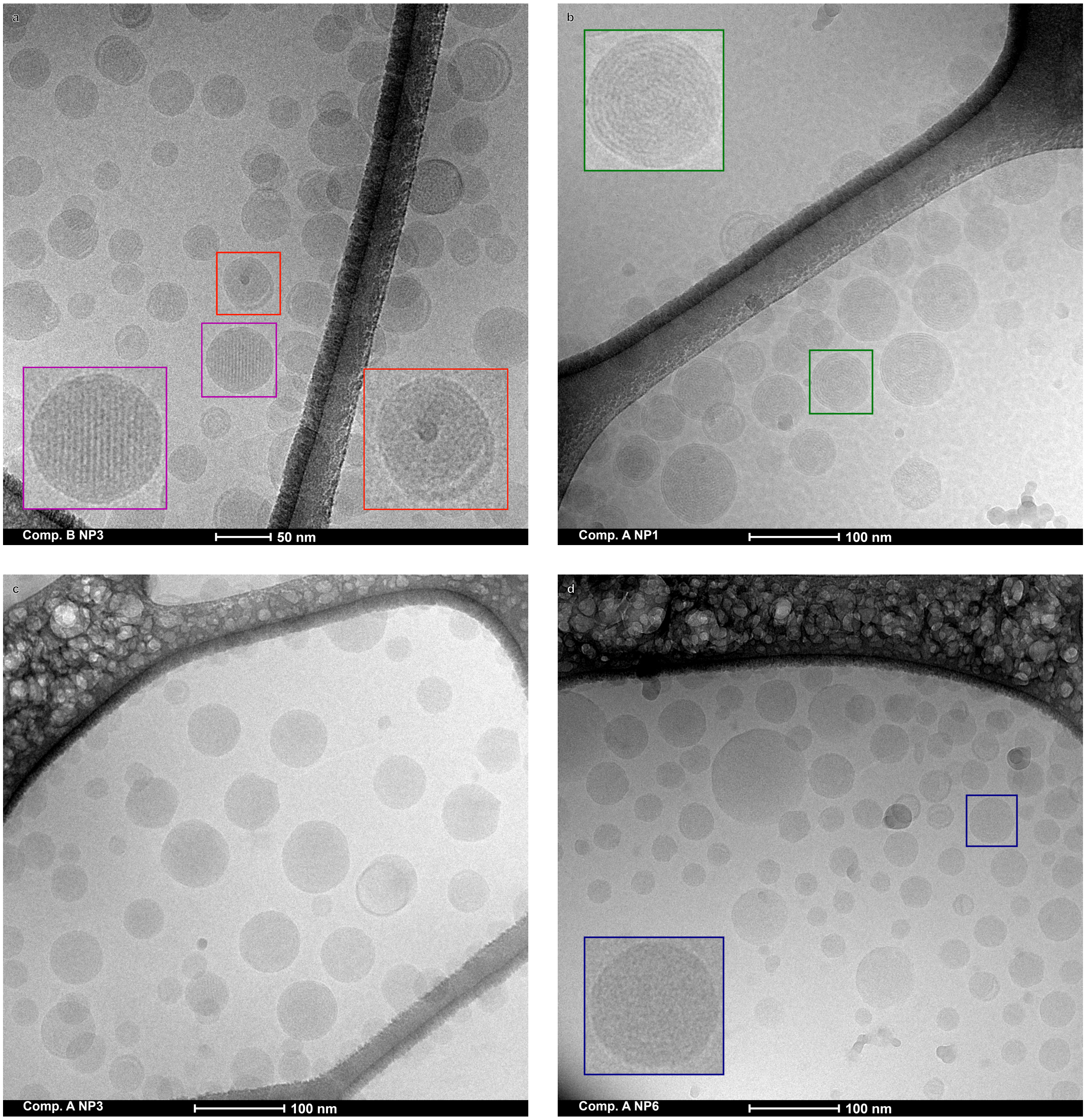


**Supplementary Figure S6: Cryo-TEM of siRNA-LNP morphologies across compositions and N/P ratios.** Representative cryogenic transmission electron micrographs of siRNA-loaded LNP formulations. **a**, Comp B (N/P 3): Coexisting multilamellar and inverse-hexagonal phases are visible; the magenta box highlights an inverse-hexagonal particle and the red box a multilamellar particle with concentric bilayer order. Amorphous and bleb-like morphologies are also present. Scale bar, 50 nm. **b**, Comp. A (N/P 1): the population is dominated by multilamellar vesicles. The green box highlights concentric bilayers consistent with siRNA intercalated between lipid layers. Scale bar, 100 nm. **c**, Comp. A (N/P 3): a heterogeneous population comprising multilamellar, amorphous and bleb-like particles. Scale bar, 100 nm. **d**, Comp. A (N/P 6):

predominantly amorphous, electron-lucent particles with occasional bleb-like surface lobules; the blue box highlights a representative low-order particle. Scale bar, 100 nm.

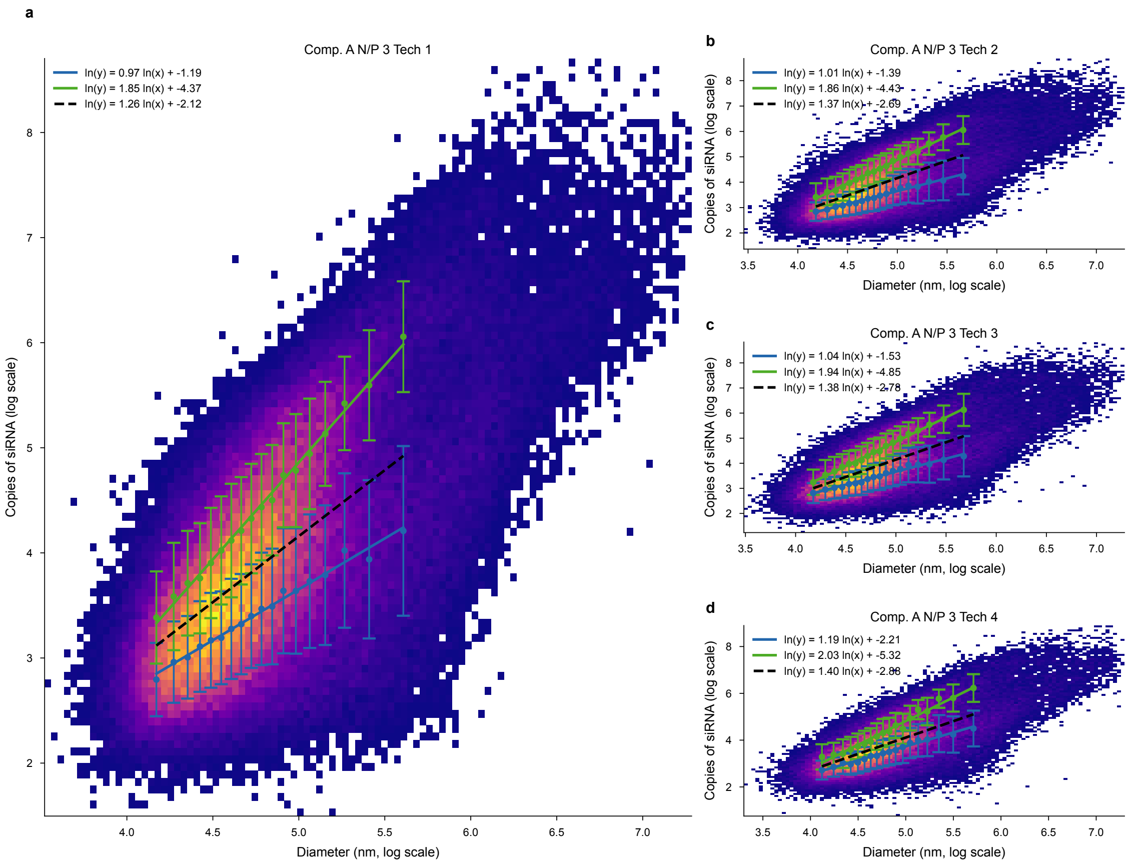


**Supplementary Figure S7: Log-log scaling of siRNA copy number with LNP radius for Composition A (N/P 3). a-d**, Two-dimensional scatter plot (log-log axes) of siRNA copy number versus LNP radius for technical replicates 1–4. Color scale (purple to yellow) indicates particle count per bin. Power-law fits are shown for the high-order subpopulation (green), the low-order subpopulation (blue) and the overall population (black dashed). a, Enlarged view of replicate 1. Green and blue circles and vertical bars denote the Gaussian mixture model component means ± s.d. within each radius bin; the legend denotes the power-law fits in the log space. Two distinct scaling regimes are resolved across all four replicates, consistent with the coexistence of high-order and low-order packing subpopulations.

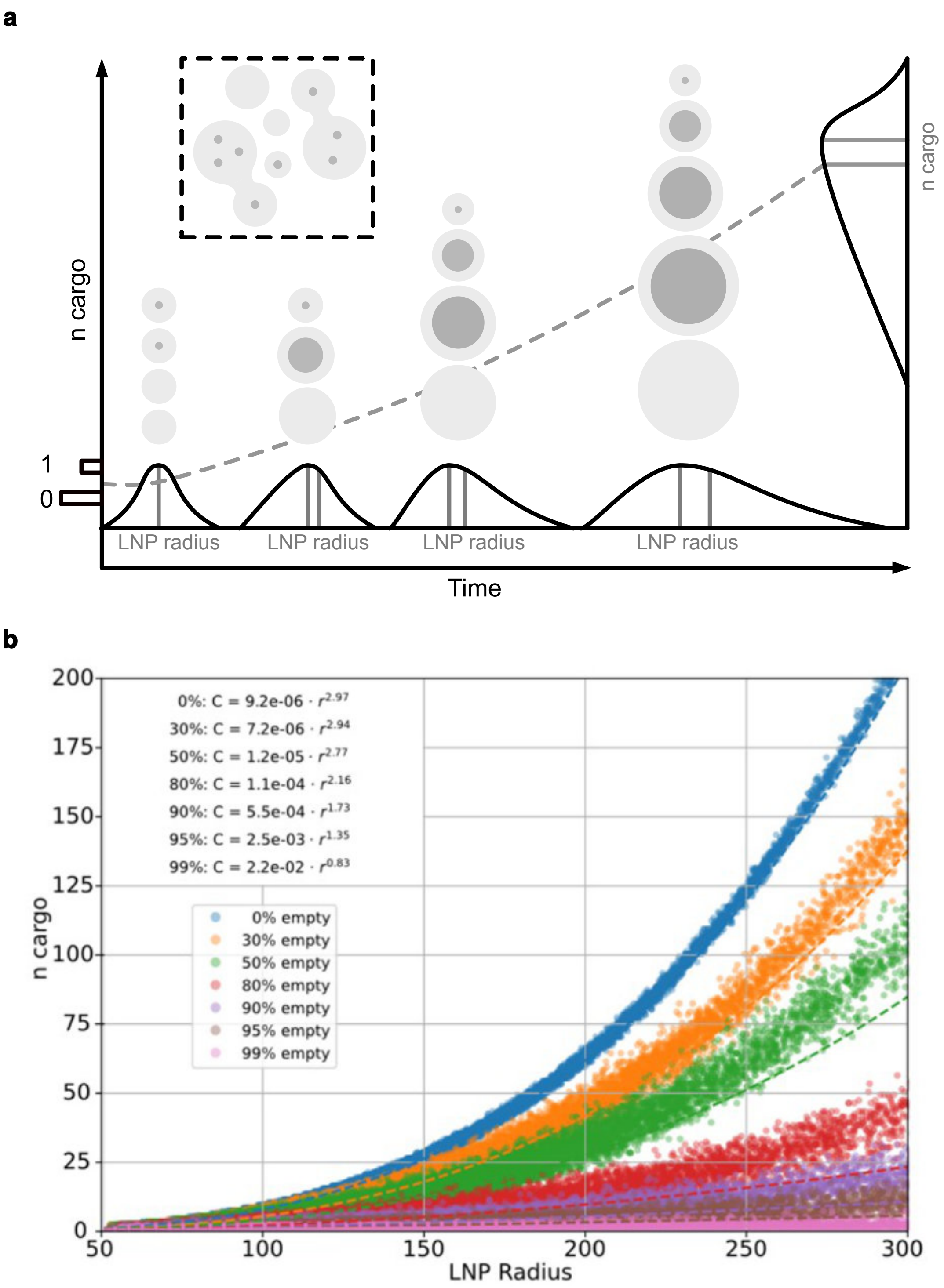


**Supplementary Figure S8: Stochastic merger model of siRNA-LNP scaling behavior. a**, Schematic of the stochastic merger of spherical volumes (SMSV) model. Precursor vesicles, cargo-loaded (dark grey) and empty (light grey), coalesce into progressively larger LNPs. The fraction of initially loaded precursors determines the cargo-size scaling relationship (dashed line). **b**, Simulated cargo copies number (*n* cargo) versus LNP radius for seven fractions of initially empty precursors (0–99%, colored as indicated in the legend). Dashed lines show power-law fits ($C = Ar^B$) with exponents annotated in the upper left. Complete initial loading (0% empty, blue) yields near-cubic scaling ($B \approx 2.97$), whereas increasing the empty fraction progressively reduces the exponent below 2.

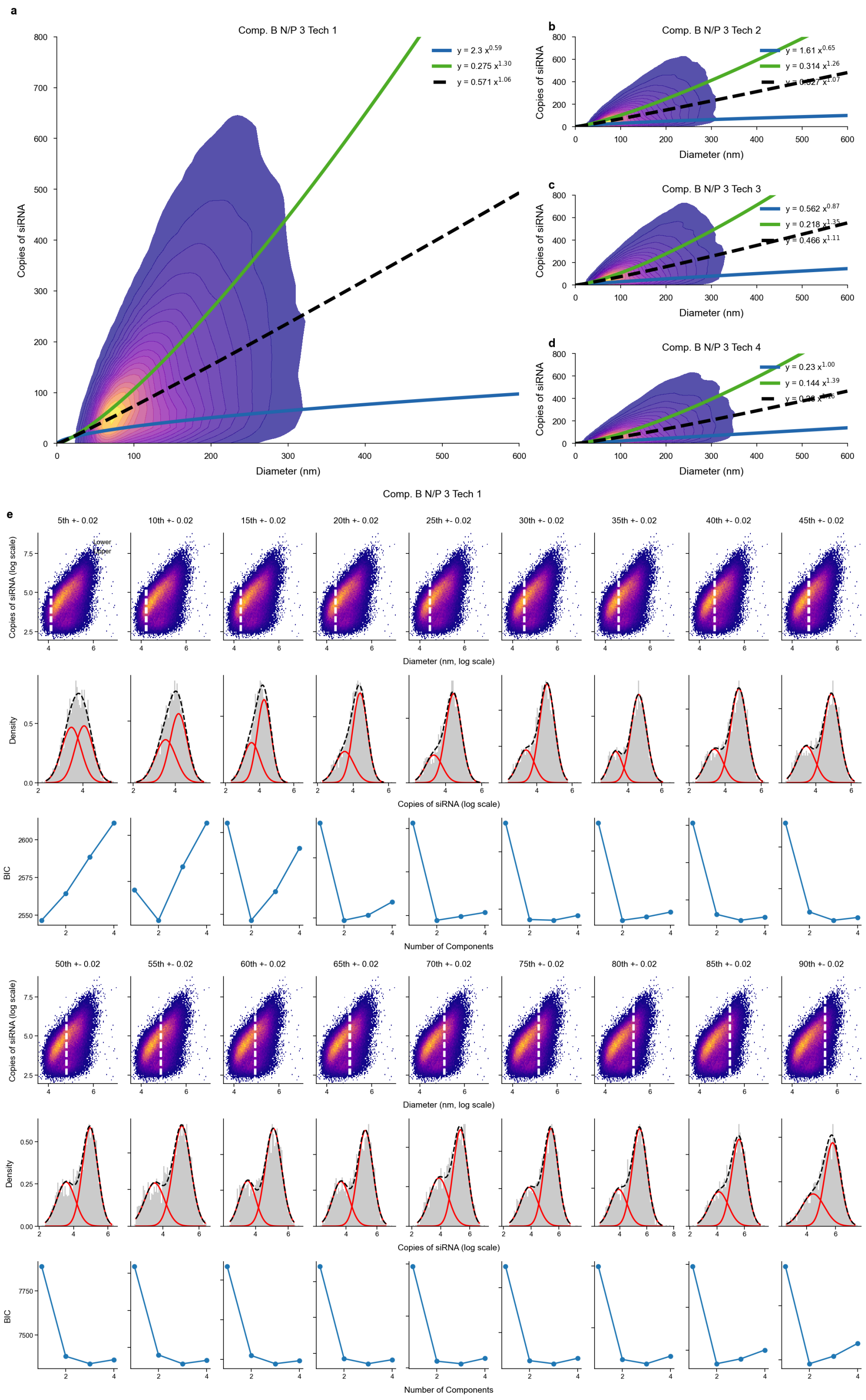


**Supplementary Figure S9: Replicate analysis of siRNA packing subpopulations for Composition B (N/P 3). a-d**, Two-dimensional kernel density estimates of siRNA copy number versus LNP diameter for four independent technical replicates (approximately 100,000–200,000 particles per replicate). Power-law fits ($y = Ax^B$) are shown for the low-order subpopulation (blue), the high-order subpopulation (green) and the overall population (black dashed). All replicates consistently resolve two distinct loading subpopulations. **e**, Gaussian mixture model (GMM) analysis of replicate 1, shown as representative, across 18 diameter percentile bins. Top row: Scatter plots of siRNA copy number versus diameter (log scale), colored by local density (purple to yellow); white dashed lines mark the percentile bin whose siRNA copy-number data are shown in the corresponding histogram below. Middle row: log-transformed siRNA copy-number distributions within each bin (grey histograms) overlaid with individual Gaussian components (red curves) and the summed fit (black curves). Bottom row: Bayesian information criterion (BIC) as a function of the number of mixture components (1–4); the minimum indicates the preferred model.

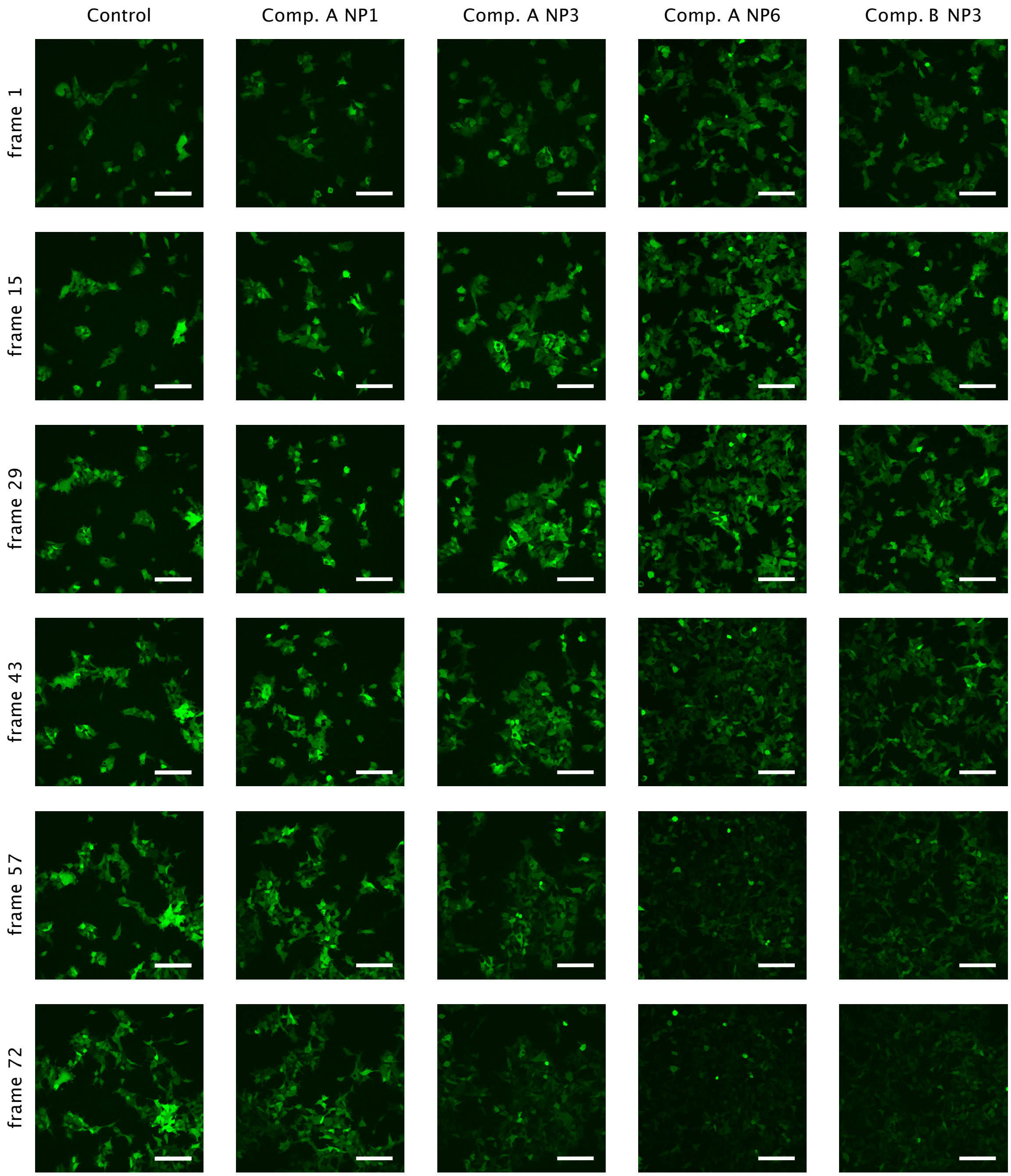


**Supplementary Figure S10: Time-lapse imaging of siRNA-mediated d2eGFP knockdown in live HEK293 cells.** Representative spinning-disk confocal fluorescence images of HEK293-d2eGFP cells treated with siRNA-loaded LNPs. Columns correspond to treatment conditions (from left to right: untreated control, Composition A at N/P 1, 3 and 6, and Composition B at N/P 3). Rows show six time points sampled from a 24-hour imaging series acquired at 20-minute intervals (frames 1, 15, 29, 43, 57 and 72, corresponding to approximately 0, 5, 9, 14, 19 and 24 hours after treatment). The maximum-intensity z-plane for eGFP fluorescence (green) is displayed; brightness and contrast settings are identical across all panels. Scale bars, 50 μm.

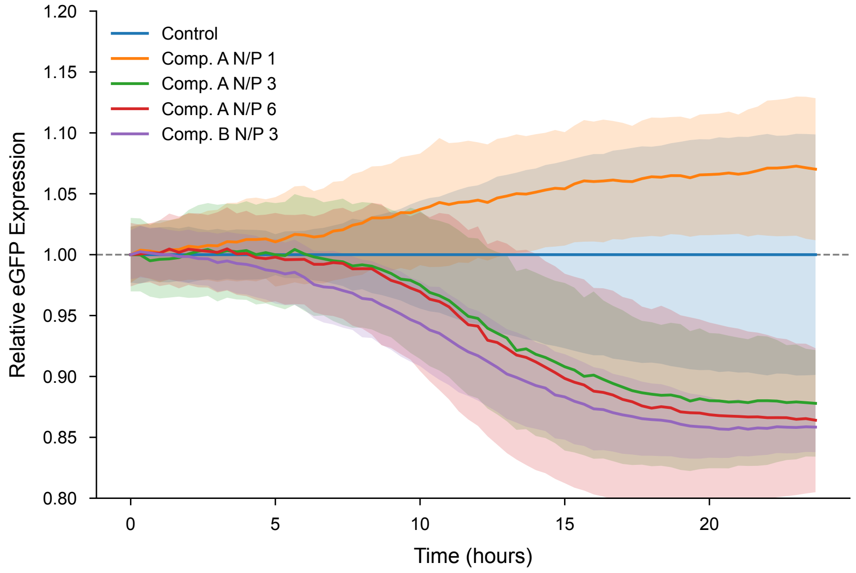


**Supplementary Figure S11: Normalized d2eGFP fluorescence trajectories during siRNA-mediated knockdown.** Control-normalized mean eGFP fluorescence intensity over 24 hours (Δt = 20 min) for HEK293-d2eGFP cells treated with siRNA-loaded LNPs. Solid lines show mean trajectories for the untreated control (blue, n = 4 replicates), Composition A at N/P 1 (orange), N/P 3 (green) and N/P 6 (red), and Composition B at N/P 3 (purple) (n = 3 replicates each). Shaded envelopes indicate ± 1 s.d. across replicates. All trajectories are normalized to the pooled control at each time point and scaled to unity at $t_0$, isolating relative silencing dynamics from differences in starting fluorescence. The horizontal dashed grey line marks the control baseline (relative intensity = 1).

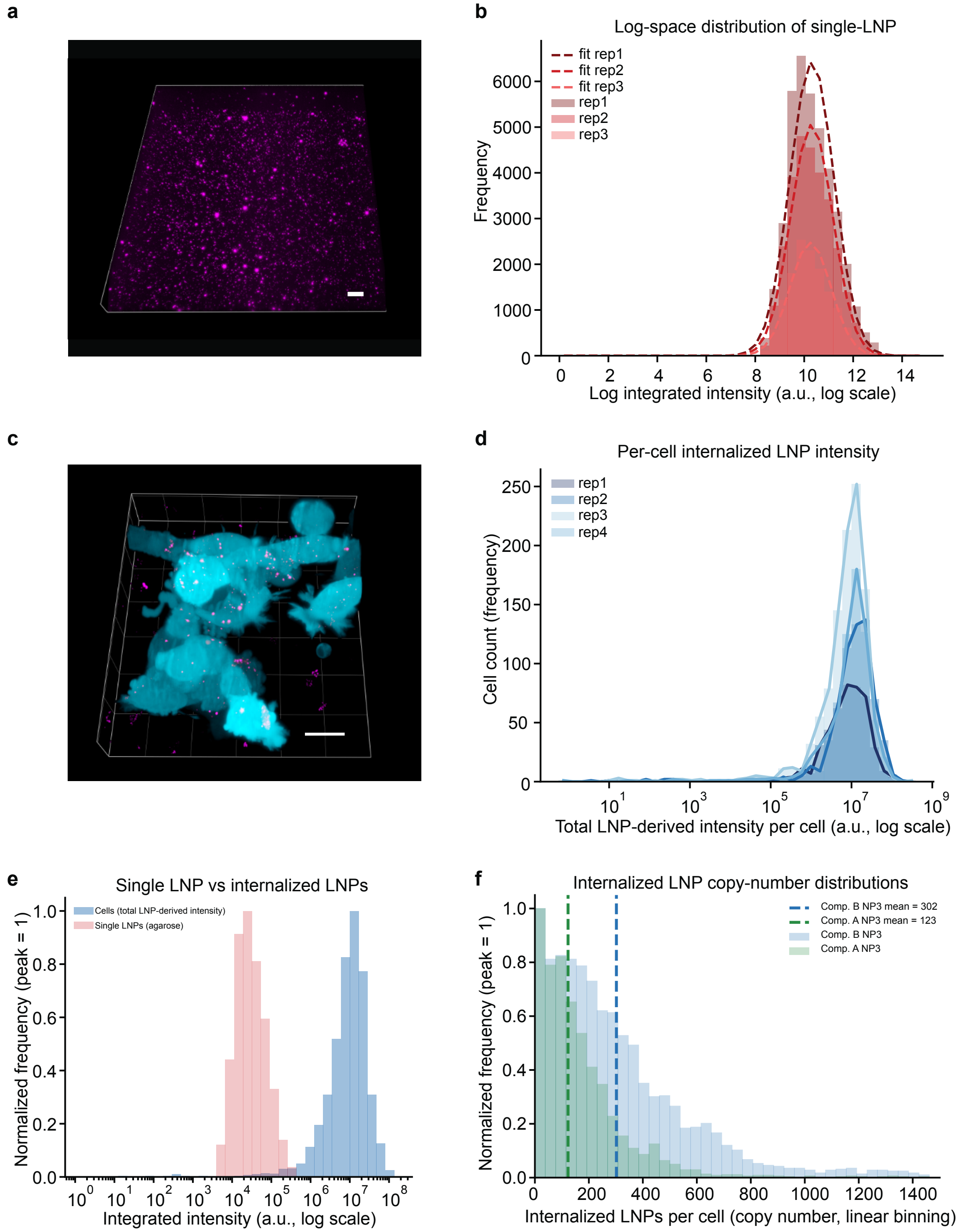


**Supplementary Figure S12: Lattice light-sheet quantification of internalized LNP copy number.** **a**, Representative three-dimensional lattice light-sheet microscopy (LLSM) volume rendering of Composition B (N/P 3) LNPs (magenta) immobilized in agarose, used for point spread function calibration and single-particle intensity extraction. Scale bar, 20 µm. **b**, Log-transformed integrated fluorescence intensity distributions of single LNPs from three technical replicates (histograms, dark to light red); dashed curves show Gaussian fits in log space used to derive calibration parameters. **c**, Volumetric LLSM rendering of d2eGFP-HEK293 cells (cyan) after 150 min incubation with

Composition B (N/P 3) LNPs (magenta). Three-dimensional cell segmentation masks were derived from the eGFP signal. Scale bar, 20 μm. **d**, Distributions of total background-corrected LNP fluorescence intensity per cell for four biological replicates (dark to light blue), plotted on a logarithmic intensity axis. **e**, Comparison of single-particle intensity (pink, from agarose calibration) and per-cell internalized LNP intensity (blue, from live-cell experiments), both peak-normalized. The cellular distribution is shifted to higher values and broadened, consistent with internalization of multiple LNPs per cell. **f**, Internalized LNP copy-number distributions for Composition B (N/P 3, blue) and Composition A (N/P 3, green), obtained by dividing each cell's integrated LNP signal by the mean single-particle brightness. Dashed vertical lines indicate mean copy numbers (302±37 and 123±4, respectively). Composition B (N/P 3) shows higher uptake than Composition A (N/P 3) under identical conditions.

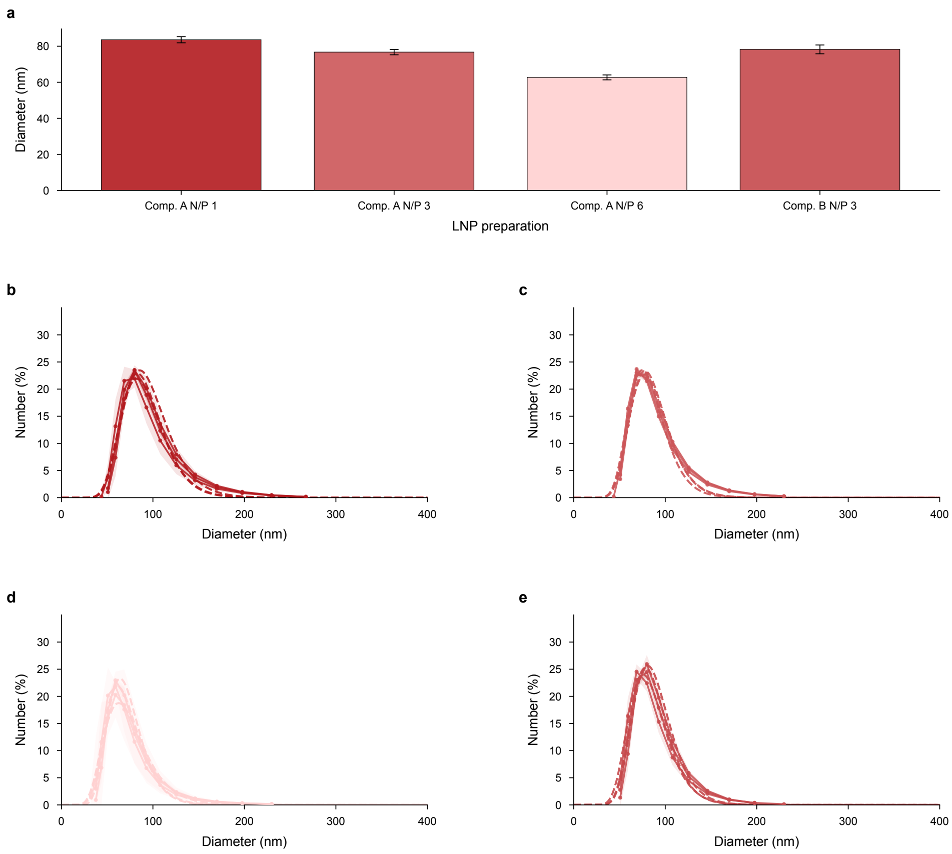


**Supplementary Figure 13: DLS characterization of LNP formulations used in live-cell experiments. a**, Mode hydrodynamic diameters of the four LNP formulations used for live-cell knockdown assays: Composition A at N/P 1, 3 and 6, and Composition B at N/P 3. Bars represent the mode diameter from number-weighted size distributions; error bars denote s.d. (n = 3–4 technical replicates). Bar shading scales with mode diameter from light pink (Composition A N/P 6, smallest) to dark red (Composition A N/P 1, largest). **b–e**, Number-weighted size distributions for Composition A at N/P 1 (b), N/P 3 (c) and N/P 6 (d), and Composition B at N/P 3 (e), respectively. Individual replicates are shown as dashed lines; the shaded envelope indicates ± 1 s.d. across replicates. All formulations display monomodal distributions with mode diameters spanning approximately 63–84 nm.

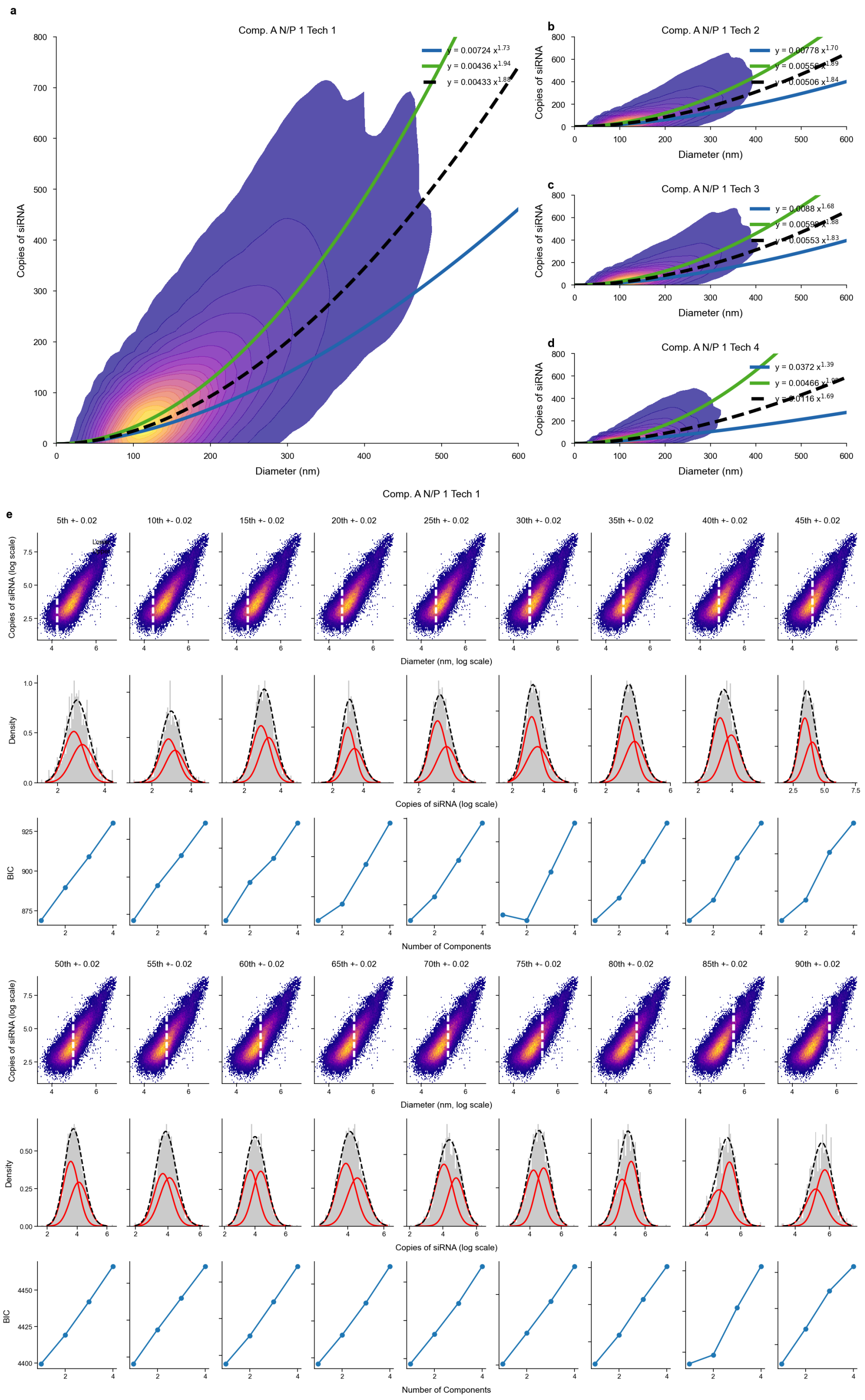


**Supplementary Figure S14: Replicate analysis of siRNA packing subpopulations for Composition A (N/P 1). a-d**, Two-dimensional kernel density estimates of siRNA copy number versus LNP diameter for four independent technical replicates (approximately 100,000–200,000 particles per replicate). Power-law fits ($y = Ax^B$) are shown for the low-order subpopulation (blue), the high-order subpopulation (green) and the overall population (black dashed). All replicates consistently resolve two distinct loading subpopulations. **e**, Gaussian mixture model (GMM) analysis of replicate 1, shown as representative, across 18 diameter percentile bins. Top row: Scatter plots of siRNA copy number versus diameter (log scale), colored by local density (purple to yellow); white dashed lines mark the percentile bin whose siRNA copy-number data are shown in the corresponding histogram below. Middle row: log-transformed siRNA copy-number distributions within each bin (grey histograms) overlaid with individual Gaussian components (red curves) and the summed fit (black curves). Bottom row: Bayesian information criterion (BIC) as a function of the number of mixture components (1–4); the minimum indicates the preferred model.

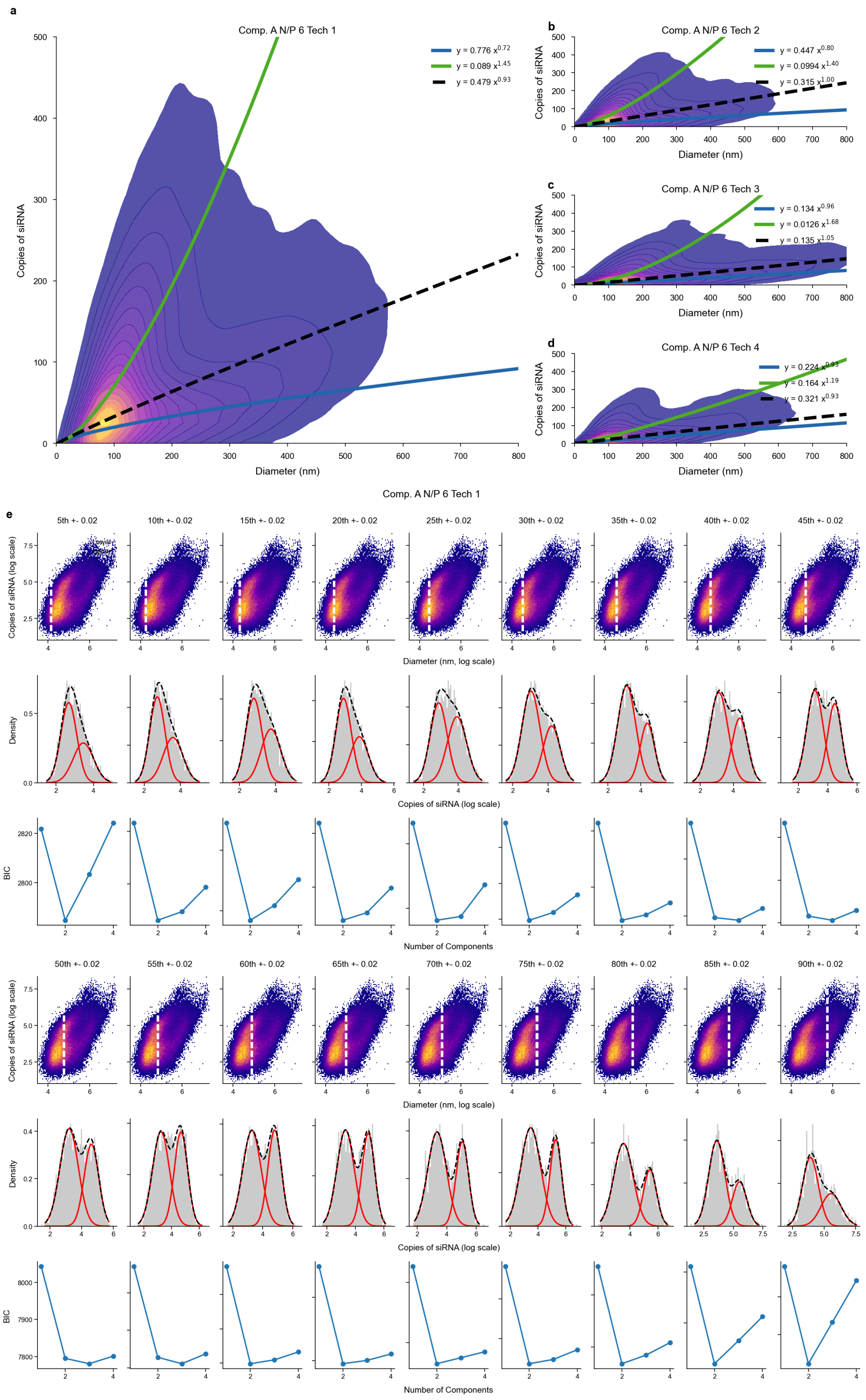


**Supplementary Figure S15: Replicate analysis of siRNA packing subpopulations for Composition A (N/P 6). a-d**, Two-dimensional kernel density estimates of siRNA copy number versus LNP diameter for four independent technical replicates (approximately 100,000–200,000 particles per replicate). Power-law fits ($y = Ax^B$) are shown for the low-order subpopulation (blue), the high-order subpopulation (green) and the overall population (black dashed). All replicates consistently resolve two distinct loading subpopulations. **e**, Gaussian mixture model (GMM) analysis of replicate 1, shown as representative, across 18 diameter percentile bins. Top row: Scatter plots of siRNA copy number versus diameter (log scale), colored by local density (purple to yellow); white dashed lines mark the percentile bin whose siRNA copy-number data are shown in the corresponding histogram below. Middle row: log-transformed siRNA copy-number distributions within each bin (grey histograms) overlaid with individual Gaussian components (red curves) and the summed fit (black curves). Bottom row: Bayesian information criterion (BIC) as a function of the number of mixture components (1–4); the minimum indicates the preferred model.
